# Engineering of human myotubes toward a mature metabolic and contractile phenotype

**DOI:** 10.1101/2023.06.12.544344

**Authors:** Simon I. Dreher, Paul Grubba, Christine von Toerne, Alessia Moruzzi, Jennifer Maurer, Thomas Goj, Andreas L. Birkenfeld, Andreas Peter, Peter Loskill, Stefanie M. Hauck, Cora Weigert

**Affiliations:** Institute for Clinical Chemistry and Pathobiochemistry, Department for Diagnostic Laboratory Medicine, University Hospital Tübingen, 72076 Tübingen, Germany; Metabolomics and Proteomics Core Helmholtz Center Munich, German Research Center for Environmental Health, 85764 Neuherberg, Germany; NMI Natural and Medical Sciences Institute at the University of Tübingen, 72770 Reutlingen, Germany; Department for Microphysiological Systems, Institute of Biomedical Engineering, Faculty of Medicine, Eberhard Karls University Tübingen, 72074 Tübingen, Germany; German Center for Diabetes Research (DZD), 85784 Neuherberg, Germany; Institute for Diabetes Research and Metabolic Diseases of the Helmholtz Zentrum München, University of Tübingen, 72076 Tübingen, Germany; Department of Internal Medicine IV, University Hospital Tübingen, 72076 Tübingen, Germany

## Abstract

1.

Skeletal muscle mediates the beneficial effects of exercise, thereby improving insulin sensitivity and reducing the risk for type 2 diabetes. Current human skeletal muscle models *in vitro* are incapable of fully recapitulating its physiological functions especially muscle contractility. By supplementation of insulin-like growth factor 1 (IGF1), a growth factor secreted by myofibers in vivo, we aimed to overcome these limitations. We monitored the differentiation process starting from primary human CD56-positive myoblasts in the presence/absence of IGF1 in serum-free medium in daily collected samples for 10 days. IGF1-supported differentiation formed thicker multinucleated myotubes showing physiological contraction upon electrical pulse stimulation following day 6. Myotubes without IGF1 were almost incapable of contraction. IGF1-treatment shifted the proteome toward skeletal muscle-specific proteins that contribute to myofibril and sarcomere assembly, striated muscle contraction, and ATP production. Elevated *PPARGC1A*, MYH7 and reduced MYH1/2 suggest a more oxidative phenotype further demonstrated by higher abundance of proteins of the respiratory chain and elevated mitochondrial respiration. IGF1-treatment also upregulated GLUT4 and increased insulin-dependent glucose uptake compared to myotubes differentiated without IGF1.

To conclude, utilizing IGF1, we engineered human myotubes that recapitulate the physiological traits of skeletal muscle *in vivo* superior to established protocols and overcome limitations of previous standards. This novel “easy to use” model enables investigation of exercise on a molecular level.

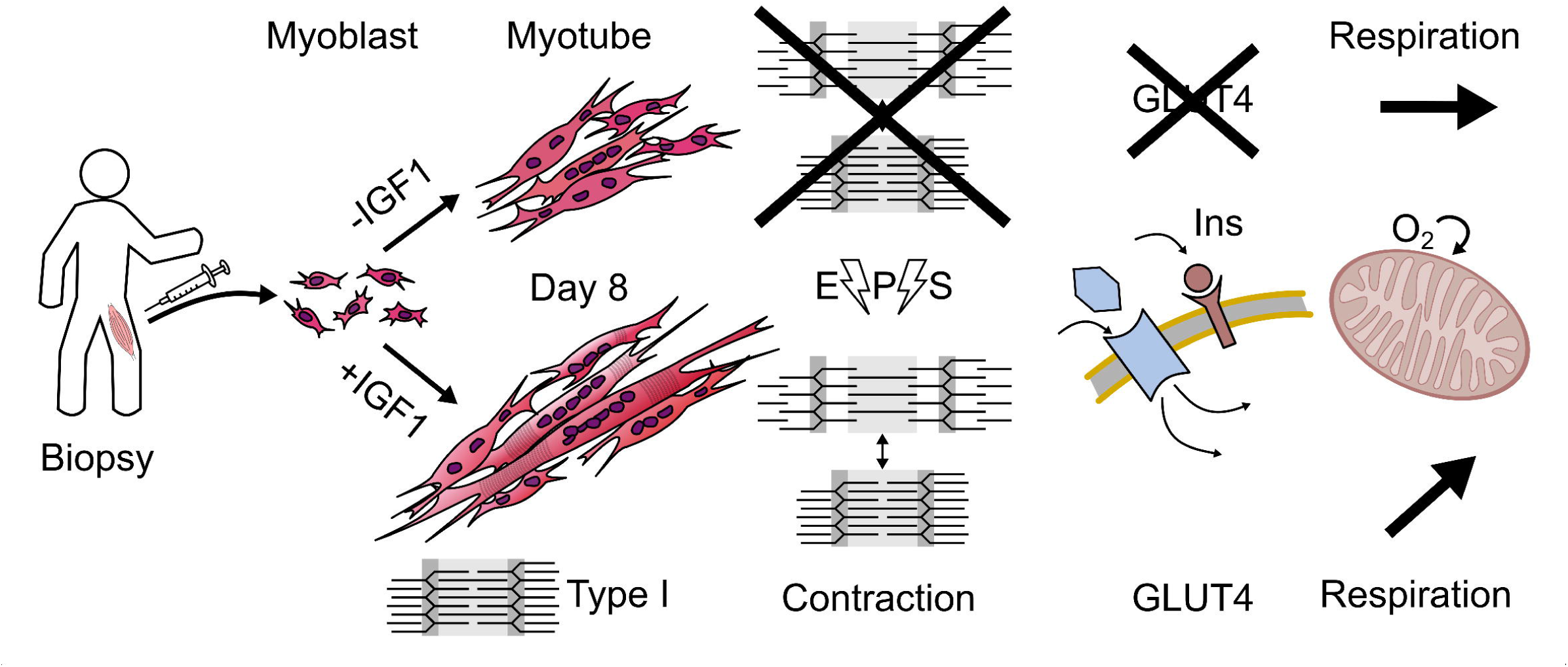

## 2. Introduction

Increased physical activity is a major part of most lifestyle intervention programs as it can preserve insulin sensitivity in healthy persons and restore compromised insulin sensitivity in subjects with prediabetes and diabetes [1–3]. As the predominantly utilized organ, skeletal muscle and its relevance for the beneficial effects of physical activity have been extensively characterized [4–6]. Adaptations involved in this metabolic benefit are increased skeletal muscle mass, capillarization, mitochondrial content and respiration thereby supporting skeletal muscle glucose disposal in response to insulin [4; 7; 8]. Skeletal muscle also represents the organ responsible for over 80% of insulin-dependent glucose uptake in the human body [5; 9]. In addition, it contributes to systemic adaptive processes by releasing myokines [10] and myometabokines to regulate signaling pathways involved in exercise adaptation in an auto-, para- or endocrine manner [11]. However, our understanding of the communication between the exercising muscle and different organs and cell types is still incomplete. This is particularly true for the systemic, health-promoting effects of exercise [12]. To study biological processes such as exercise on a whole-body level and its multi-organ response and interconnection on a molecular level requires physiologically relevant models. To answer many research questions, screening the impact of compounds or metabolites on a system and sampling of tissue biopsies of internal organs cannot be performed in humans directly which leaves researchers with two options. Common mouse models, while allowing to study the interplay between organ systems come with inherent ethical limitations and, most importantly, limited transferability of findings to the human system due to metabolic and anatomical differences [13; 14]. Alternatively, and close to the system of interest, *in vitro* models of primary human cells are used and their interplay in co-cultures or organ-on-chip models have gained interest in recent years especially in light of the first of the 3Rs (Replacement, Reduction and Refinement) to replace animal models [15; 16].

*In vitro* protocols for differentiation of myotubes from primary human satellite cells have existed for decades [17–19] and have been further optimized over the years from using rat or human brain extract [20; 21] over 2% serum (horse or fetal calf) [22; 23] to serum-free protocols [24; 25]. They consistently yield long, fused, multinucleated myotubes that recapitulate some aspects of myofibers *in vivo* and are a widely used tool to answer specific molecular questions. Nevertheless, the current *in vitro* differentiation models are yet incapable of fully recapitulating the physiological functions of myofibers *in vivo* especially regarding muscle contraction and key features of its metabolic function e.g. mitochondrial respiration and glycemic control [26]. Human myotubes barely contract during electrical-pulse-stimulation (EPS) [27; 28], they predominantly produce ATP by glycolysis, rather than mitochondrial oxidative phosphorylation [29], the insulin-responsive glucose uptake is reduced and the ratio of GLUT4 to GLUT1 expression is low compared to adult skeletal muscle [30; 31]. During stem cell and precursor cell differentiation, growth factors are needed to guide proper differentiation. The growth factor insulin-like growth factor 1 (IGF1) is secreted by myofibers during muscle movement *in vivo* and is responsible for hypertrophy, and production of muscle-specific and contractile apparatus proteins [32–34]. Additionally, IGF1 already could have played a small part in previous promising yet complicated multi-step differentiation attempts to overcome some of the functional limitations of primary human myotubes *in vitro* [25].

Aim of the study was to utilize IGF1 to engineer a more physiological myotube model from primary human precursor cells *in vitro*. We used a serum-free differentiation protocol to differentiate myotubes from primary human myoblasts supplemented with or without 100 ng/ml IGF1 corresponding to total IGF1 concentrations in human plasma [35; 36]. Over a time course of 10 days, we took daily samples to characterize differences between the two differentiation protocols in the proteome and on transcript level, used immunohistochemistry to visualize cell fusion and structural proteins, applied immunoblotting and glucose uptake assay to study the insulin responsiveness, measured mitochondrial respiration using Seahorse and assessed the capability to functionally contract by EPS and video analysis.

## 3. Materials and Methods

### 3.1. Cell Culture

Primary human myoblasts were obtained from muscle biopsies as described previously [37]. Biopsy donors were participants of two previous exercise intervention studies [38; 39] and only baseline biopsies were used. For the time course experiment (Figure 1), biopsies from a total of n=4 donors (2 female, 2 male, age 36±12 (24-52), BMI 34±5 (27.62-39.98)) were used and in total n=14 donors (10 female, 4 male, age 33±9 (21-52), BMI 30±4 (23.61-39.98)) for the additional analysis of mitochondrial respiration, insulin signalling, glucose uptake, and immunofluorescence staining. Myoblast isolation and enrichment of CD56+ myoblasts by magnetic bead cell sorting are described in [24]. Cell culture surfaces were prepared with a non-gelling thin-layer GelTrex™ (Thermo Fisher Scientific, Germany) coating. Myoblasts were proliferated in cloning media (α-MEM:Ham’s F-12 (1:1), 20% (v/v) FBS, 1% (v/v) chicken extract, 2 mM L-glutamine, 100 units/ml penicillin, 100 µg/ml streptomycin, 0.5 µg/ml amphotericin B) until 90% confluency. Myotube differentiation was induced on day 0 and maintained for 10 days in fusion media (α-MEM, 2 mM L-glutamine, 50 µM palmitate, 50 µM oleate (complexed to BSA with a final BSA concentration of 1.6 mg/ml in medium), 100 µM carnitine) with or without supplementation of 100 ng/ml (13.16 nmol/l) IGF1 (human recombinant IGF1, I3769, Sigma-Aldrich, Germany). Medium was changed three times per week and 48 hours before harvest. For the time course-experiment, cells were harvested on day 1 of myotube differentiation and every day from day 3 to day 10 (Figure 1).

**Figure 1.**
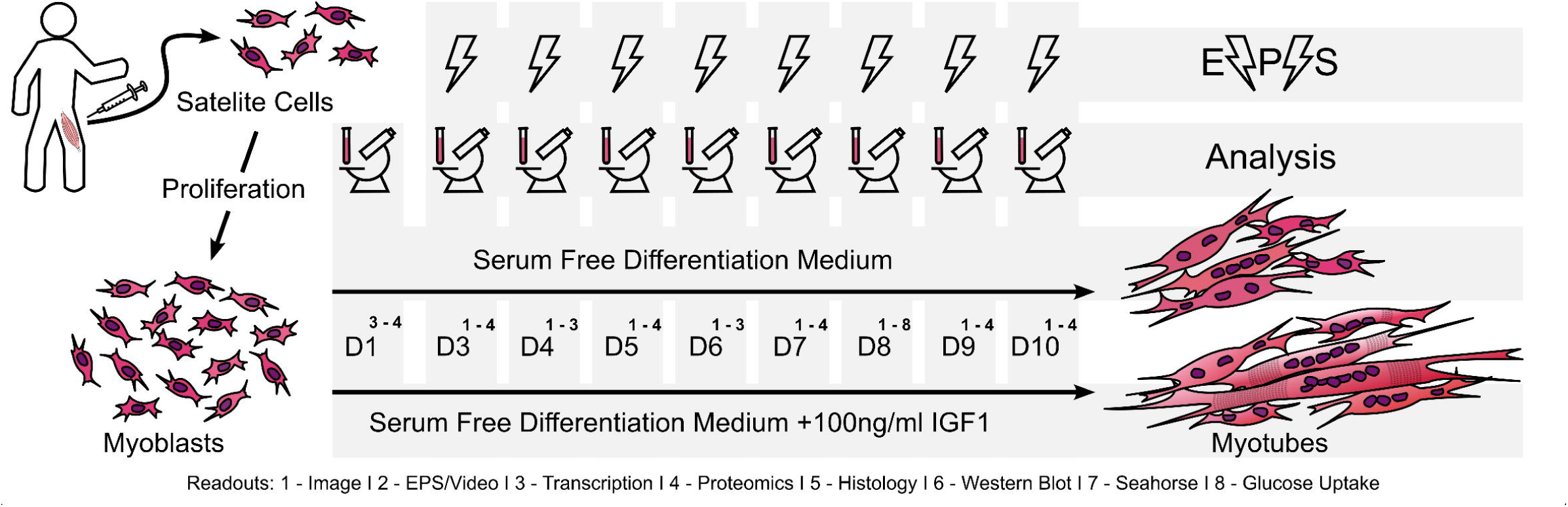
Experimental design. Primary human satellite cells isolated from vastus lateralis muscle biopsies were proliferated to myoblasts and differentiated into myotubes for 10 days in serum-free differentiation medium in the presence or absence of 100ng/ml IGF1. On day 1 and daily from day 3 myotubes were sampled and analyzed. Cells were imaged, electrical-pulse-stimulation (EPS) was performed and transcription was analyzed daily from day 3 to day 10. From samples obtained on day 1 proteomic analysis were performed and transcription was analyzed. From samples obtained on days 3, 5 and 7 to 10 proteomic analysis were performed. Functional analysis including immunostaining, phospho-Western-blotting, respiratory measurement by Seahorse and glucose uptake were conducted on day 8.

### 3.2. Electrical-Pulse-Stimulation

For electrical-pulse-stimulation (EPS), cells were cultured in compatible 6-well plates (Falcon, Corning, USA) in parallel. From day 3 of myotube differentiation, EPS was performed for 4h at 1Hz, 2ms, 25V using C-Pace EP (Ionoptix, USA) and 3 videos of 15 s at 60 fps per group and donor were taken in random spots of each well using an Axiovert 40C (Carl Zeiss Microscopy, Germany) with Flexacam C3 Camera (Leica, Germany). To achieve tetanic contraction, EPS was performed at 10Hz, 2ms, 25V for 5sec followed by a resting phase.

### 3.3. Video Analysis and Movement Index

Myotube contraction speed, directionality and adherence to EPS frequency were analysed using the open heart ware (OHW) software [40]. A total of 189 randomly taken videos, 3 videos per group and timepoint over the time course from day 4 to day 10 were evaluated by 6 independent blinded observers that evaluated amount of movement based on reference videos with a value between 0 and 3. Value 0 corresponds to no movement, 1: slight contraction/movement of a partial structure/myotube, 2: clear contraction/movement of a partial structure/myotube or slight contraction/movement of a complete structure/myotube 3: clear contraction/movement of a complete structure/myotube. For each timepoint, condition and donor, a single movement index between 0 and 9 was calculated by first calculating the mean value per video over all observers and subsequently calculating the sum of all three videos per timepoint, condition and donor. Movement index of all n=4 donors were used for statistical evaluation.

### 3.4. Proteomics

#### Mass spectrometry (MS) sample preparation

Cells were lysed in RIPA buffer (25 mM Tris pH7.6, 150 mM NaCl, 3.5 mM SDS, 12.1 mM sodium deoxycholate, 1% v/v Triton X100) and incubated at 4 °C for 30 min. Lysates were then subjected to ice-cold sonication bath for 30 s and another incubation (4 °C, 15 min). The protein concentration of the individual samples was determined using BCA assay following the instruction manual (Pierce Biotechnology, USA). BSA was used as an internal standard. Ten µg of sample were enzymatically proteolysed using a modified filter-aided sample preparation (FASP) protocol as described in [41; 42]. Peptides were stored at −20 °C until MS measurement.

#### MS measurements

LC-MSMS analysis was performed in data-dependent acquisition (DDA) mode. MS data were acquired on a Q-Exactive HF-X mass spectrometer (Thermo Fisher Scientific, Germany) coupled online to a nano-RSLC (Ultimate 3000 RSLC; Dionex). Tryptic peptides were automatically loaded on a C18 trap column (300 µm inner diameter (ID) × 5 mm, Acclaim PepMap100 C18, 5 µm, 100 Å, LC Packings) at 30 µl/min flow rate. For chromatography, a C18 reversed phase analytical column (nanoEase MZ HSS T3 Column, 100 Å, 1.8 µm, 75 µm x 250 mm, Waters, USA) at 250 nl/min flow rate in a 95 minutes non-linear acetonitrile gradient from 3 to 40 % in 0.1 % formic acid was used. The high-resolution (60.000 full width at half-maximum) MS spectrum was acquired with a mass range from 300 to 1.500 m/z with automatic gain control target set to 3 x 10^6^ and a maximum of 30 ms injection time. From the MS prescan, the 15 most abundant peptide ions were selected for fragmentation (MSMS) if at least doubly charged, with a dynamic exclusion of 30 seconds. MSMS spectra were recorded at 15.000 resolution with automatic gain control target set to 5 x 10^2^ and a maximum of 50 ms injection time. The normalized collision energy was 28, and the spectra were recorded in profile mode.

#### MS data processing and protein identification

Proteome Discoverer software (Thermo Fisher Scientific; version 2.5) was used for peptide and protein identification via a database search (Sequest HT search engine) against Swissprot human data base (Release 2020_02, 20349 sequences), considering full tryptic specificity, allowing for up to one missed tryptic cleavage site, precursor mass tolerance set to 10 ppm, fragment mass tolerance to 0.02 Da. Carbamidomethylation of cysteine was set as a static modification. Dynamic modifications included deamidation of asparagine and glutamine, oxidation of methionine; and a combination of methionine loss with acetylation on protein N-terminus. Percolator was used for validating peptide spectrum matches and peptides, accepting only the top-scoring hit for each spectrum, and satisfying the cut-off values for FDR <1 %, and posterior error probability <0.05. The final list of proteins complied with the strict parsimony principle. Protein identification with a Sequest HT score of below 1.6 were excluded from further analysis.

#### MS label-free quantification

The quantification of proteins was based on the area under the curve of the abundance values for unique peptides. Abundance values were normalized against total abundance to account for sample loading errors. The protein abundances were calculated averaging the abundance values for up to 50 admissible peptides. The final protein ratio was calculated as median of all peptide comparisons between all samples. The statistical significance of the ratio change was ascertained employing the T-test approach described [43], which is based on the presumption that expression changes are screened for proteins that are just a few in comparison to the number of total proteins being quantified. The quantification variability of the non-changing “background” proteins can be used to infer which proteins change their expression in a statistically significant manner.

To ensure robustness of results, a change in protein abundance was considered significant with a Benjamini-Hochberg (BH) corrected p-value below 0.05 only if detected in at least three out of four donors in at least one experimental group.

### 3.5. Gene Expression

RNA was extracted employing the RNeasy kit (Qiagen, Germany). For mRNA detection, reverse transcription was carried out with the Transcriptor First Strand Synthesis Kit (Roche, Switzerland). Expression was measured using QuantiFast SYBR Green PCR Mix and QuantiTect Primer Assays (*MYH1*: QT01671005; *MYH2*: QT00082495; *MYH7*: QT00000602; *PPARGC1A* (*PGC1α*): QT00095578; *GLUT1*: QT00068957; *GLUT4*: QT00097902; *RPS28*: QT02310203; *TBP*: QT00000721; Qiagen, Germany) in a LightCycler 480 (Roche, Switzerland). mRNA standards for PCR were generated by purifying PCR-product (MinE-lute PCR Purification Kit, Qiagen, Germany) and 10-fold serial dilution.

### 3.6. Immunofluorescence staining

Myotubes cultured on 1-well cell culture chamber on PCA slide (94.6140.102 Sarsted, Germany) were fixed on day 8 with ROTI-Histofix 4% (Roth, Germany) for 20 min at room temperature before storage at 4°C. Fixed myotubes were blocked and permeabilized in 3% (m/m)-BSA solution with 0.1% (m/m) Digitonin (D5628, Sigma-Aldrich, Germany) in DPBS (Thermo Fisher Scientific, Germany) for 30 min at room temperature. Proteins were stained with monoclonal anti-Myosin, MHC-fast, Clone MY-32 (1:2000, M4276, Sigma-Aldrich, Germany) and monoclonal anti-Myosin, MHC-slow, Clone NOQ7.5.4D (1:4000, M8421, Sigma-Aldrich, Germany) primary antibodies at 4°C overnight, Goat anti-Mouse IgG1 Cross-Adsorbed Secondary Antibody, Alexa Fluo 488 (1:2000, A21121, Invitrogen, USA) secondary antibody, DAPI (1:500, Invitrogen, USA) and Rhodamine Phalloidin (1:300, Invitrogen, USA) for 2 h at room temperature and imaged using the ApoTome System (Carl Zeiss Microscopy, Germany).

### 3.7. Mitochondrial respiration

Myoblasts were cultured directly on the Seahorse assay plates (Seahorse XFe24 FluxPak, Agilent, USA) with 20,000 cells/well. On day 8 of myotube differentiation, medium was removed and cells were washed according to manufacturer’s protocol with Seahorse assay medium consisting of DMEM (103575-100, Agilent, USA) with 10 mM glucose, 2 mM L-glutamine, 1 mM pyruvate. After calibration of the Seahorse XFe24 Analyzer (Agilent, USA), respiration was measured using freshly prepared substrates in Seahorse assay medium: 10 µM oligomycin (Port 1), 20 mM FCCP (Port 2), and 5 µM rotenone and 5 µM antimycin A (Port 3). After measurement, cells were lysed in RIPA buffer (25 mM Tris pH7.6, 150 mM NaCl, 3.5 mM SDS, 12.1 mM sodium deoxycholate, 1% (v/v) Triton X100) and total protein content determined in 10 µl lysate by BCA assay (Pierce Biotechnology, USA) in duplicates. Oxygen consumption rate (OCR) and extracellular acidification rate (ECAR) were normalized to protein content. Seahorse measurement was performed in quadruplicates of n=4 individual donors. The calculation of mitochondrial respiration was carried out according to manufacturer’s protocol using Wave software. Total oxygen consumption (OCR) is the area under the curve (AUC) for the total measurement. Basal respiration is baseline OCR (no injection) minus non-mitochondrial respiration (OCR after Rotenone and Antimycin A injection). Proton leak is OCR after Oligomycin injection minus non-mitochondrial respiration. ATP production is the difference between basal respiration and proton leak. Uncoupled (maximal) respiration is OCR after FCCP injection minus non-mitochondrial respiration. Spare capacity is the difference between uncoupled respiration and basal respiration. The energy map was drawn for each measurement by plotting OCR/ECAR and mean±SD after calculation of the quotient.

### 3.8. Immunoblotting

Protein lysates were prepared in RIPA buffer (25 mM Tris pH7.6, 150 mM NaCl, 3.5 mM SDS, 12.1 mM sodium deoxycholate, 1% (v/v) Triton X100) containing cOmplete EDTA-free protease inhibitor (Roche Diagnostics, Switzerland) and phosphatase inhibitor (1 mM NaF, 0.5 mM sodium pyrophosphate, 1 mM β-glycerophosphate, 1 mM sodium orthovanadate). Protein concentration was determined by BCA assay (Pierce Biotechnology, USA) and samples heated in Laemmli sample buffer for 5 minutes to 95°C. Sodium dodecyl sulfate polyacrylamide (7.5–15%) gradient gel electrophoresis and semidry electroblotting (transfer buffer: 48 mM Tris, 39 mM glycine, 1.3 mM SDS, and 20% (v/v) methanol) were performed using nitrocellulose membranes (Whatman, UK). NET buffer (150 mM NaCl, 50 mM Tris/HCl, pH 7.4, 5 mM EDTA, 0.05% (v/v) Triton X-100, and 0.25% (m/m) gelatin) was used for blocking. Proteins were detected using GLUT4 (1:2000, PA5-23052, Invitrogen, USA), AKT(1:500, 610860, BD, USA), pAKT(Ser-473) (1:1000, 9271L, cell signaling, USA) and GAPDH (1:20000, ab8245, abcam, USA) primary antibodies, overnight at 4°C, and IRDye 680RD Goat anti-Mouse IgG Secondary Antibody (1:20000, 926-68070, Li-COR, USA) and IRDye® 800CW Goat anti-Rabbit IgG Secondary Antibody (1:10000, 926-32211, Li-COR, USA) for 2 hours at room temperature. Signals were detected on an Odyssey scanner (Li-COR, USA), quantification of signal intensity was performed with ImageStudio. GLUT4 protein levels were normalized on GAPDH, pAKT referred to total AKT.

### 3.9. Glucose Uptake

Glucose uptake was determined using the Glucose Uptake-Glo™ Assay (Promega, USA) following the protocol provided by the manufacturer. Myotubes grown on 24 well plates were fasted for 3 h in glucose-free DMEM (103575-100, Agilent, USA). Incubation with 100 nM insulin (Roche Diagnostics, Switzerland) was performed for one hour at 37 °C followed by one-hour incubation at room temperature after adding 10 mM 2-deoxyglucose. Luminescence was measured using the GloMax® Multi Detection System (Promega, USA) after one hour incubation at room temperature with the provided detection reagent. Obtained relative light units (RLU) were donor normalized.

### 3.10. Data analysis and statistics

Linear regression models were analyzed with R4.1.1/RStudio [44]. Normality was tested by Shapiro-Wilk-test from the R package ‘stats’ (v3.6.3) and non-normal data were log-transformed. For correction of multiple testing p-values were adjusted with Benjamini-Hochberg FDR<5% (BH). Plotted protein abundances based on proteomic analysis reflect median values over all detected peptides. Data were given as mean±SD and BH corrected p-values <0.05 were considered significant. Differences between all timepoints were assessed using one-way ANOVA with Fisheŕs LSD post hoc test or Bonferroni correction for multiple comparisons when appropriate. Graphs were made using the R packages ‘ggplot2’ (v3.3.2) and ‘ggrepel’ (v0.9.1) and figures assembled using InkScape (v1.0). Functional enrichment analysis was carried out online using https://biit.cs.ut.ee/gprofiler/ with p<0.05 as threshold and g:SCS as method for multiple testing correction. Homo sapiens was chosen as organism and analysis was performed using the databases from gene ontology biological process (GO:BP), cellular components (GO:CC), molecular functions (GO:MF), Reactome (REAC), KEGG, Wikipathways (WP) and human protein atlas (HPA). Ingenuity pathway analysis (IPA) for upstream regulators was performed based on proteomics datasets comparing +IGF vs −IGF.

## 4. Results

### 4.1. Improved Differentiation and Contraction in +IGF Myotubes

Microscopic evaluation of differentiating myotubes revealed optical differences between myotubes differentiated with (+IGF) or without IGF1 (−IGF) starting around day 4 and 5 (Figure S1). While both protocols yielded elongated fused myotubes, from day 5-6 thicker myotubes with bigger diameters were visible in the presence of IGF1. From day 6-7 these myotubes showed a more pronounced three-dimensional phenotype compared to the myotubes fused without IGF1 (Figure S1). On day 8, a clear striated pattern representing the contractile apparatus was only visible in the presence of IGF1 (Figure 2 A). When we applied electrical-pulse-stimulation (EPS) to myotubes differentiated without IGF1 typically no contraction was visible (Figure 2 B-D). The heatmap of moving areas generated by the OHW software shows only areas outside of cells that represent background and air bubbles moving (Figure 2 C), which is also represented by the basically flat line representing movement speed over time (Figure 2 D). In contrast, the myotubes differentiated with IGF1 showed pronounced contraction (Figure 2 E, F) that nicely adhered to the 1Hz provided by EPS (Figure 2 G). The movement curve (Figure 2 G) demonstrates controlled and fast contraction from relaxed resting position 1 to fully contracted at 3 with the most movement at 2 towards full contraction at 3 and a slower relaxation after the end of the pulse from 3 over the most movement towards relaxation at 4 to the next relaxed resting position (Figure 2 G). We quantified muscular contraction by calculating the movement index, screening 189 videos, 3 videos per group and timepoint randomly taken (Figure 2 H). Contraction started to be visible around day 5 of differentiation with IGF1 increasing in frequency and intensity until day 8 and 9 where physiological contraction is visible in all donors differentiated with IGF1 while cellular movement in response to EPS was almost completely absent in myotubes differentiated without IGF1 (Figure 2 H). For reference, we provide videos representing best, average and worst contractions found in myotubes differentiated with IGF1 (Video 1-7) as well as representative videos showing myotubes differentiated without IGF1 during EPS (Video 8-10). We frequently observed spontaneous contractions only in myotubes differentiated with IGF1 (Video 11-13). In one case we managed to film spontaneous contraction followed by controlled contraction with EPS (Video 13). By increasing stimulation to 10Hz, myotubes differentiated with IGF1 can perform tetanic contractions repeatedly, demonstrating the potential of our model to functionally simulate different exercise modalities *in vitro* (Video 14). In summary, differentiation of primary human myoblasts with IGF1 supplementation leads to visibly thicker myotubes with striated pattern formation and functional contractibility induced by EPS.

**Figure 2.**
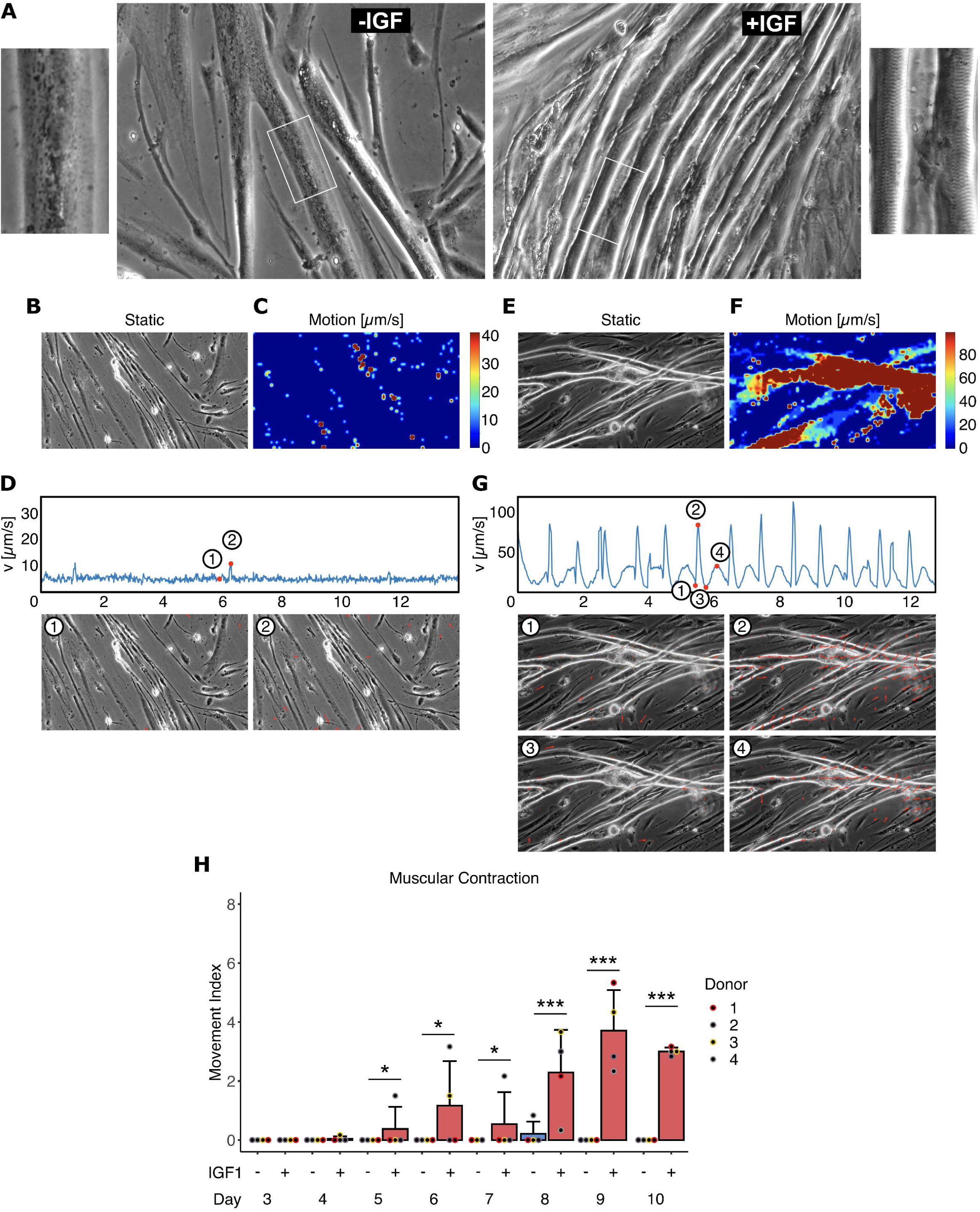
Human myotube morphology and contraction. Primary human myotubes differentiated in the presence or absence of IGF1 to analyze morphology and capability for contraction in response to EPS. A) Microscopic images of differentiated myotubes on day 8 of differentiation. B-G) Representative output of the video analysis with the open heart wave (OHW) program for myotubes differentiated without (B-D) or with IGF1 (E-G). B,E) Static frame of the video; C,F) Heatmap representing degree and area of movement; D,G) Movement over time curve with static frames and arrow plots visualizing contraction in response to EPS at 1Hz. H) Contractability by EPS was quantified by calculating the movement index (0-9) based on randomly taken videos between day 3 and day 10 in myotubes differentiated without (blue bars) or with IGF1 (red bars). Bars represent mean ± SD, individual data points are depicted. Significant differences were assessed using one-way ANOVA with Fisher’s LSD post hoc test, *p < 0.05, **p < 0.01, ***p < 0.001, n = 4 individual donors.

### 4.2. Myotubes with Altered Proteome

We used mass spectrometry-based proteomics to assess the time-dependent adaptations in the cellular proteome of the myotubes differentiated with or without IGF1. In total, 5456 proteins were detected and only considered significantly changed at a corrected p-value below 0.05 (BH) and when measured in at least three out of four donors in one experimental group (Figure 3 A). During differentiation, the abundance of a comparable number of proteins was changed without IGF1 and in the presence of IGF1. Comparing +IGF vs −IGF samples at each timepoint, between 102-165 proteins showed higher abundance and between 88-139 showed lower abundance (Figure 3 A). Principle component analysis (PCA) over all samples revealed no outliers but a clear separation comparing +IGF vs −IGF along PC 2 (Figure 3 B). Stained for group and day of differentiation, the PCA showed a clear separation along PC 1 with progressing differentiation from day 1 to day 10 as well as different trajectories for −IGF (bottom left to top right) and +IGF (bottom left to bottom right) (Figure 3 C). Interestingly, the proteome did not undergo further major changes within −IGF and +IGF groups between day 7 and day 10. On day 8, both proteomes show the greatest distance from day 1 indicating potentially fully differentiated myotubes. The proteome data suggest that the two different protocols lead to different proteomes showing that −IGF was incapable of differentiating toward the full phenotype of +IGF even if more time was given to fully differentiate (Figure 3 C). When looking at the top differentially abundant proteins comparing +IGF and −IGF on day 8 we found many proteins important for skeletal muscle ATP supply (CKMT2, PYGM, PGAM2), overall muscular development (MLIP, CSRP3, MUSTN1, MYMK, FHL3) and contractile apparatus (MYL1, MYOM2, MYOM1, MYL2, MYBPC2, MYL11, TNNC2) (Figure 3 D). All these proteins were more abundant in myotubes differentiated +IGF.

**Figure 3.**
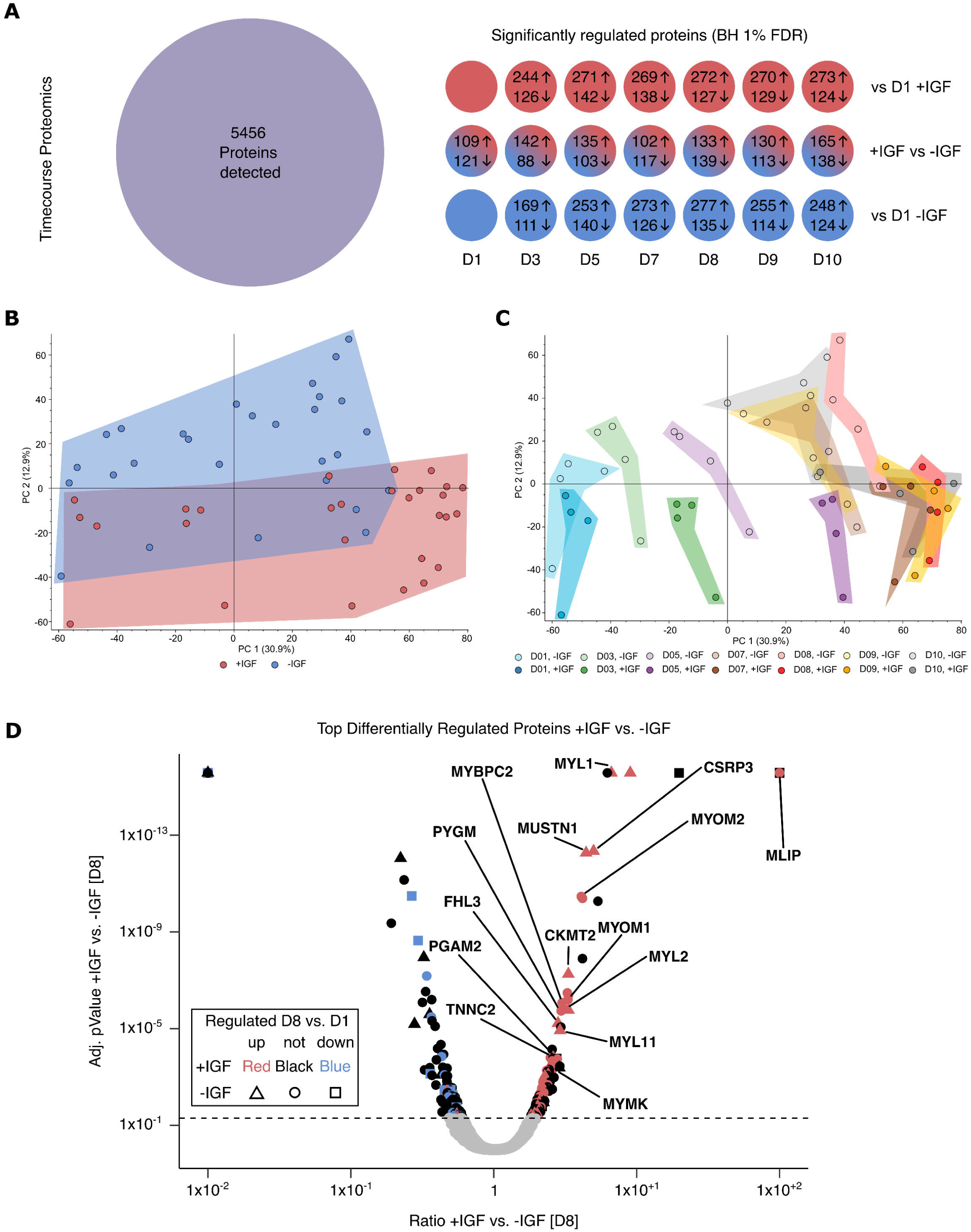
Proteomic analysis of human myotube differentiation. Primary human myotubes differentiated in the presence or absence of IGF1 were subjected to proteomic analysis on day 1, 3, 5 and 7 to 10 of differentiation. A) Number of detected and significantly regulated proteins for every timepoint comparing myotubes differentiated with (red) or without (blue) IGF1 at day 1-10 (+IGF vs −IGF) or the regulation vs. day 1 in +IGF or −IGF. B Principal component analysis (PCA) for each sample in the proteomic analysis stratified by myotubes differentiated with (red) or without (blue) IGF1. C) PCA stratified by timepoint (colors) and myotubes differentiated without (light dot and color) and with (dark dot and color) IGF1. D) Volcano-Plot depicting significantly different proteins between myotubes differentiated with or without IGF1 on day 8 of differentiation. Colors additionally indicate significant regulation vs day 1 in +IGF (red =up; blue = down) and shapes indicate significant regulation vs day 1 in −IGF (triangle =up; square = down). The color black and shape circle indicate no regulation vs day 1. Significant differences were defined by a Benjamini-Hochberg corrected p-value below 0.05, n=4 individual donors.

### 4.3. Elevated Skeletal Muscle Associated Proteins in +IGF Myotubes

To get further insight into the proteomic changes induced by IGF1 we utilized enrichment analysis on the differentially abundant proteins comparing +IGF vs −IGF at each timepoint (Figure 4). To generate an overview of the enrichment results for each day we performed a meta-analysis. We assigned each top 5 enriched term based on the up or downregulated proteins comparing +IGF vs −IGF on each day to one of the preambles “Skeletal Muscle”, “Muscle”, “Other Muscle”, “Extracellular Matrix” or “Non-Muscle”. The quotient is displayed over the time of differentiation. Enriched terms associated with muscle in general, and specifically with skeletal muscle increased continuously over time in +IGF vs −IGF while other muscle associated terms decreased during the end of differentiation between day 7-8. Skeletal muscle associated enriched proteins peak at day 8 with IGF1. Terms associated with ECM proteins were downregulated early during differentiation in +IGF and normalized at later timepoints (Figure 4 A).

**Figure 4.**
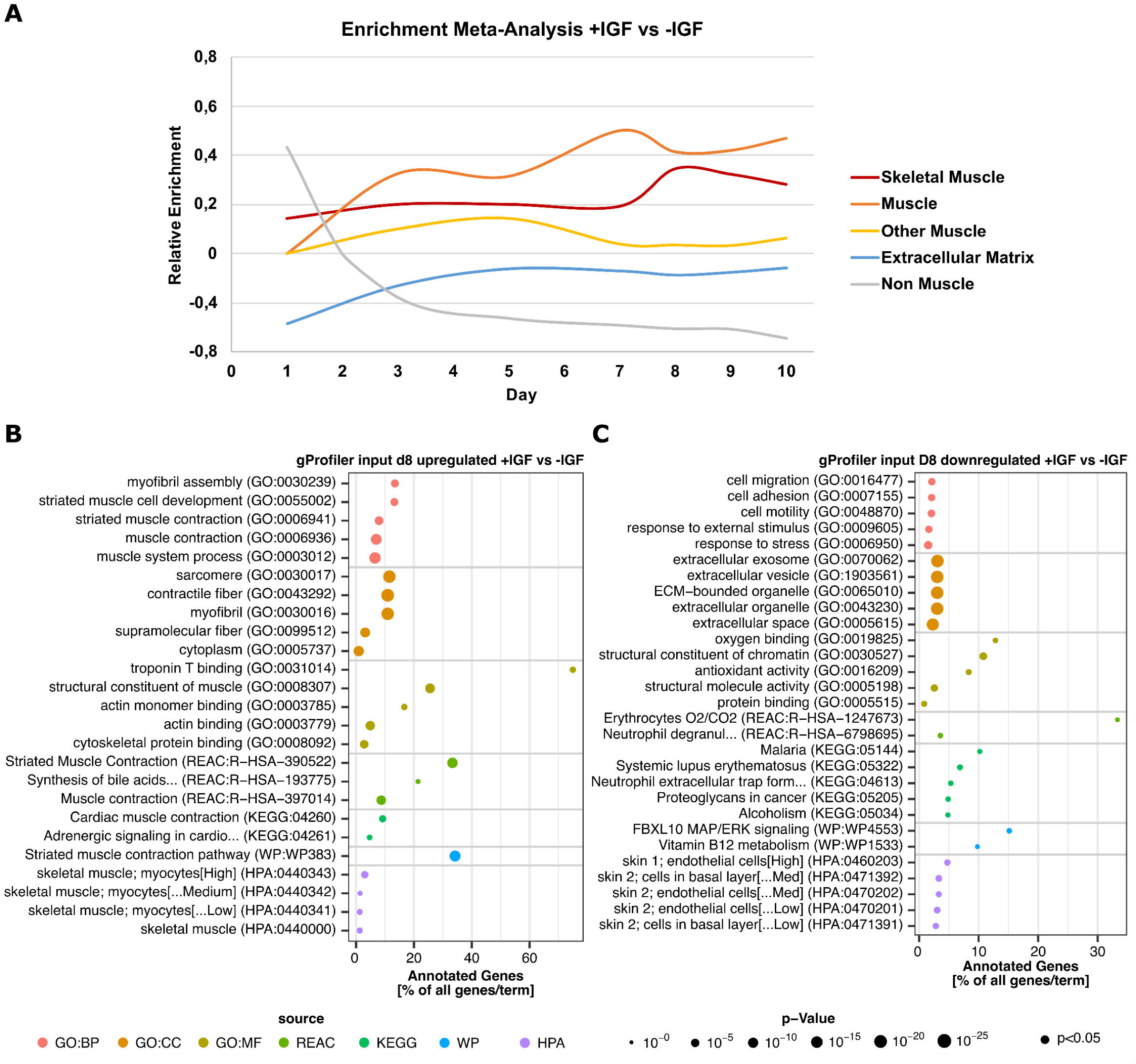
Pathway analysis during human myotube differentiation. Primary human myotubes differentiated in the presence or absence of IGF1 were subjected to proteomic analysis over 10 days of differentiation. Pathway enrichment analysis was performed comparing myotubes +IGF vs −IGF (n=4 individual donors). A) Enrichment meta-analysis, each top 5 enriched terms based on the up or downregulated proteins comparing +IGF vs −IGF on each day was assigned to one of the preambles “Skeletal Muscle”, “Muscle”, “Other Muscle”, “Extracellular Matrix” or “Non-Muscle” showing the quotient over the time of differentiation. B) gProfiler analysis based on significantly upregulated proteins on day 8 +IGF vs −IGF. C) gProfiler analysis based on significantly downregulated proteins on day 8 +IGF vs −IGF. Significant differences were defined by a Benjamini-Hochberg corrected p-value below 0.05 and detection.

On day 8, almost all enriched terms based on proteins with higher abundance in +IGF vs −IGF are associated with striated or skeletal muscle development, contractile apparatus, myofibril assembly and striated muscle contractions (Figure 4 B) while the enriched terms based on proteins with lower abundance in +IGF vs −IGF are associated with non-muscle terms or the ECM (Figure 4 C). Well in accordance, activated upstream regulators comparing +IGF vs −IGF at day 8 based on Ingenuity pathway analysis include muscle-specific transcription factors MEF2C, MYO1D and ESSRA (Figure S2 A). In summary, top regulated candidates and enrichment analysis comparing differentially abundant proteins in myotubes differentiated with or without IGF1 clearly hint toward more functional skeletal muscle differentiation and contractile apparatus assembly with IGF1. Based on the proteome data, day 8 was chosen for further functional characterization.

### 4.4. Elevated Contractile Apparatus Protein Expression in +IGF Myotubes

As our proteome analysis clearly hinted towards striated muscle and contractile apparatus (Figure 3 D, Figure 4 A-B) and optical analysis showed proper contractile apparatus assembly with striated pattern after 8 days of differentiation with IGF1 (Figure 2 A) we looked into the protein expression of the contractile apparatus proteins in more detail (Figure 5). Titin (TTN) connecting the myosins with the Z-band, the myomesins (MYOMs) connecting the myosins to form the M-band, myosin binding protein C (MYBPC1), tropomyosin (TPM1) and skeletal muscle troponins (TNNI2, TNNT3) that assemble the contractile apparatus were all elevated on protein level +IGF (Figure 5 A-H). While TTN, MYOM3, TPM1, TNNI2, and TNNT3 (Figure 5 A, D, F - H) were more upregulated +IGF, interestingly, MYOM1, MYOM2 and MYBPC1 (Figure 5 B, C, E) were only upregulated +IGF demonstrating insufficient supply of contractile apparatus proteins and thus potentially improper assembly of the contractile apparatus in the absence of IGF1.

**Figure 5.**
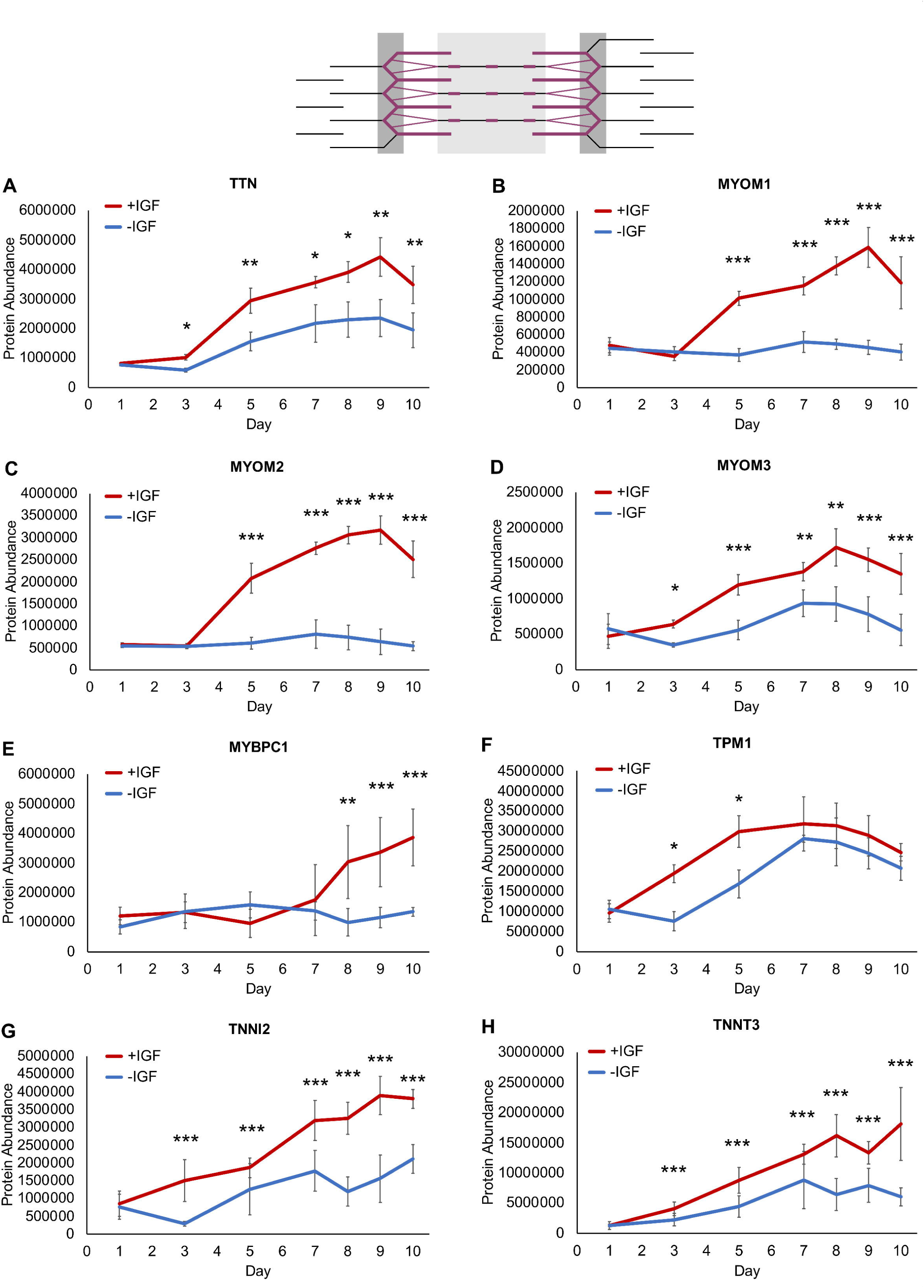
Contractile apparatus proteins during human myotube differentiation. Primary human myotubes differentiated in the presence or absence of IGF1 were subjected to proteomic analysis over 10 days of differentiation. Schematic representation of the contractile apparatus generated with InkScape (v1.0) highlights the location of analyzed proteins in purple. Regulation of contractile apparatus proteins A) TTN, B) MYOM1, C) MYOM2, D) MYOM3, E) MYBPC1, F) TPM1, G) TNNI2, H) TNNT3 was analyzed on days 1, 3, 5, 7-10 comparing protein levels between myotubes differentiated with or without IGF1. Curves represent mean ± SD, based on median values over all detected peptides. Significant differences were defined by a Benjamini-Hochberg corrected p-value below 0.05, *p < 0.05, **p < 0.01, ***p < 0.001, n = 4 individual donors.

### 4.5. Similar Fast Type II Glycolytic Fiber Marker Proteins

A key part of the contractile apparatus, also distinguishing the glycolytic from the oxidative fiber type in skeletal muscle, are the myosin heavy chains. We first looked at the fast glycolytic markers MYH1 and MYH2 and found intense staining in both myotubes differentiated without and with IGF1 on day 8 (Figure 6 A). While sometimes a slight striated pattern was seen with the MyHfast antibody (directed against MYH1 and MYH2) in myotubes differentiated without IGF1, intracellular staining was still mostly diffuse. On the contrary, a clearly structured striated pattern was found with the MyHfast staining in myotubes differentiated with IGF1 (Figure 6 A). Next, we looked at the fast type II marker MYH1, MYH2 and MYH4 RNA and protein expression (Figure 6 B-G). While RNA expression of MYH1 and MYH2 showed lower levels in myotubes +IGF, protein abundance of MYH1 and MYH2 was rather comparable in +IGF vs −IGF (Figure 6 B-E), similar to the immunohistochemistry staining (Figure 6 A). *MYH4* RNA levels were slightly elevated +IGF1 on day 9 and day 10 while MYH4 protein was less abundant in myotubes with IGF1 at the end of differentiation (Figure 6 D).

**Figure 6.**
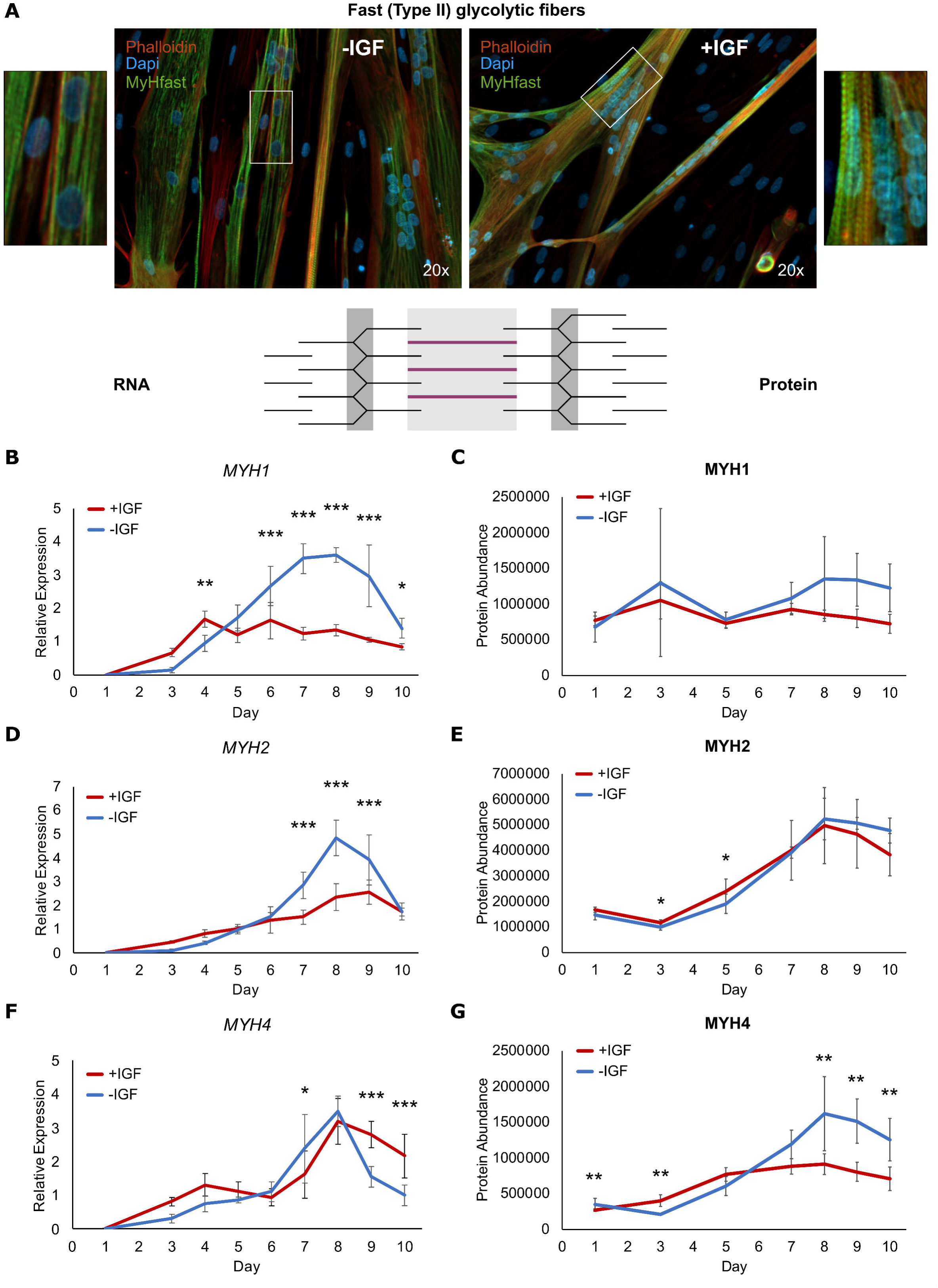
Fast (type II) glycolytic fiber markers during human myotube differentiation. Primary human myotubes differentiated in the presence or absence of IGF1 were subjected to transcriptional and proteomic analysis over 10 days of differentiation. Schematic representation of the contractile apparatus generated with InkScape (v1.0) highlights the location of analyzed proteins in purple. A) Fixed slides of myotubes differentiated with or without IGF1 on day 8 of differentiation were stained against a myosin heavy chain fast (MyHfast, directed against MYH1 and MYH2) antibody (green), phalloidin (red) and DAPI (blue). Regulation of fast (type II) glycolytic fiber markers B) *MYH1*, D) *MYH2*, F) *MYH4* RNA expression was analyzed on days 1-10 comparing expression between myotubes differentiated with or without IGF1. Curves represent mean ± SD. Significant differences were assessed using one-way ANOVA with Fisher’s LSD post hoc test, *p < 0.05, **p < 0.01, ***p < 0.001, n = 4 individual donors. Regulation of fast (type II) glycolytic fiber marker proteins C) MYH1, E) MYH2, G) MYH4 was analyzed on days 1,3, 5, 7-10 comparing protein levels between myotubes differentiated with or without IGF1. Curves represent mean ± SD, based on median values over all detected peptides. Significant differences were defined by a Benjamini-Hochberg corrected p-value below 0.05, *p < 0.05, **p < 0.01, ***p < 0.001, n = 4 individual donors.

### 4.6. More Slow Type I Oxidative Fiber Marker Proteins in +IGF Myotubes

Next, we studied the slow oxidative fiber type markers. In myotubes differentiated without IGF1, we found almost no MyHslow positive staining (directed against MYH7) on day 8 indicating low abundance of MYH7 while myotubes differentiated with IGF1 showed intense positive staining and again a clear striated pattern (Figure 7 A). In line with the immunohistochemical staining, RNA and protein expression of the most prominent oxidative type I marker MYH7 was stronger expressed throughout differentiation in myotubes differentiated with IGF1 (Figure 7 B-C). Additional slow oxidative fiber type markers, MYH6 and MYL3 were also elevated on RNA and protein level especially later during differentiation (day 5-10) (Figure 7 D-G). In summary, contractile apparatus proteins are more abundant in myotubes differentiated with IGF1 and histological assessment indicates proper assembly. While fast glycolytic type II fiber type markers are present with both differentiation protocols, slow oxidative type II markers are only expressed or elevated in myotubes differentiated in the presence of IGF1.

**Figure 7.**
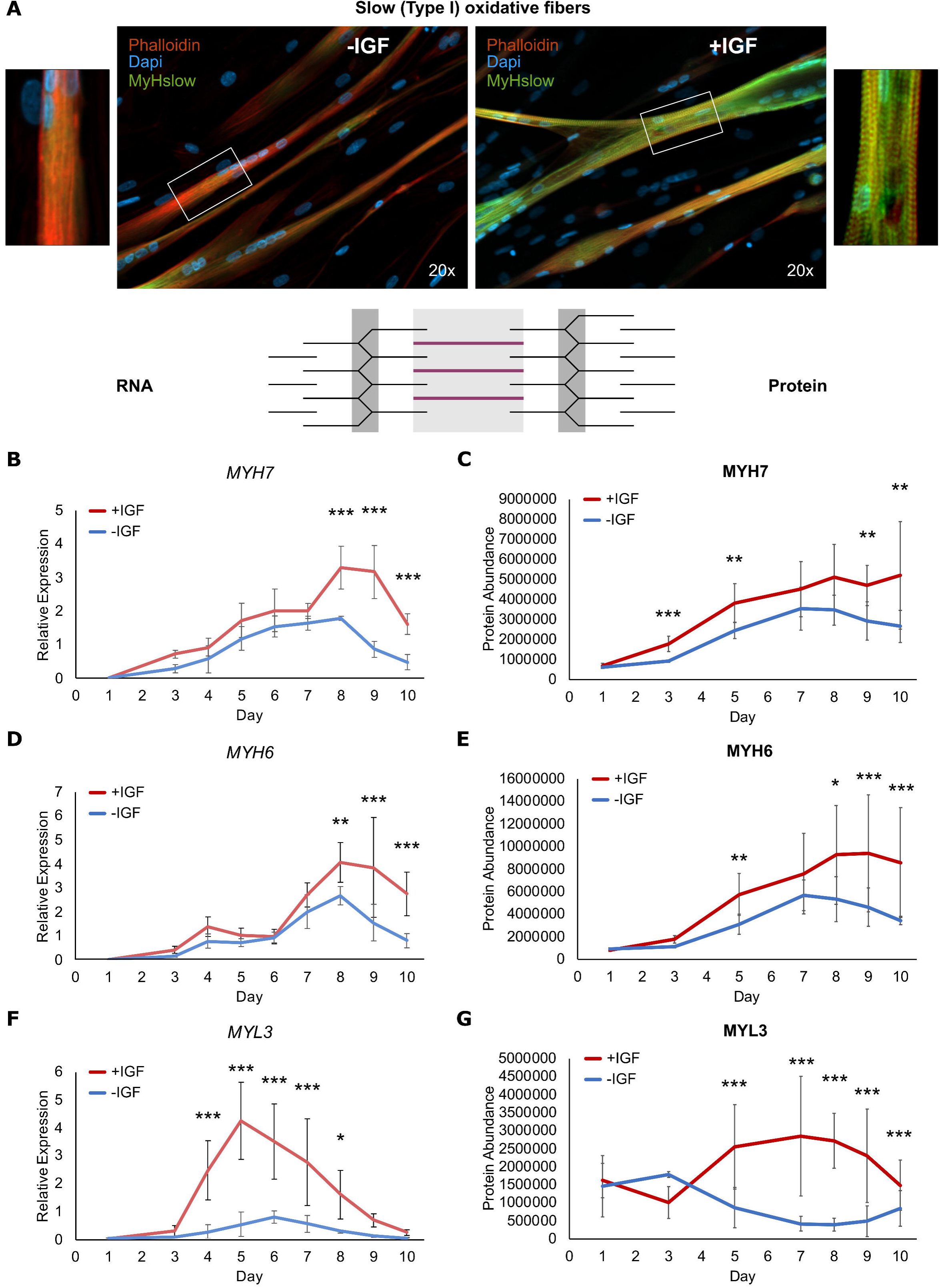
Slow (type I) oxidative fiber markers during human myotube differentiation. Primary human myotubes differentiated in the presence or absence of IGF1 were subjected to transcriptional and proteomic analysis over 10 days of differentiation. Schematic representation of the contractile apparatus generated with InkScape (v1.0) highlights the location of analyzed proteins in purple. A) Fixed slides of myotubes differentiated with or without IGF1 on day 8 of differentiation were stained against a myosin heavy chain slow (MyHslow, directed against MYH7) antibody (green), phalloidin (red) and DAPI (blue). Regulation of slow (type I) oxidative fiber markers B) *MYH7*, D) *MYH6*, F) *MYL3* RNA expression was analyzed on days 1-10 comparing expression between myotubes differentiated with or without IGF1. Curves represent mean ± SD. Significant differences were assessed using one-way ANOVA with Fisher’s LSD post hoc test, *p < 0.05, **p < 0.01, ***p < 0.001, n = 4 individual donors. Regulation of slow (type I) oxidative fiber marker proteins C) MYH7, E) MYH6, G) MYL3 was analyzed on days 1,3, 5, 7-10 comparing protein levels between myotubes differentiated with or without IGF1. Curves represent mean ± SD, based on median values over all detected peptides. Significant differences were defined by a Benjamini-Hochberg corrected p-value below 0.05, *p < 0.05, **p < 0.01, ***p < 0.001, n = 4 individual donors.

### 4.7. Elevated Energy Metabolism and Sarcoplasmic Ca^2+^ Release Proteins in +IGF Myotubes

As we found enhanced contractility and expression of contractile apparatus proteins in +IGF myotubes we next looked at proteins supporting energy supply during muscle contraction. Both striated muscle specific creatine kinases, M-type (CKM) and mitochondrial 2 (CKMT2) catalyzing phosphate transfer from creatine phosphate to ATP were significantly more abundant starting at day 3-5 in myotubes differentiated with IGF1 (Figure 8 A). Enzymes involved in glycogenolysis and glycolysis, namely glycogen phosphorylase (PYGM) and the rate limiting glycolysis enzyme phosphofructokinase (PFKM) as well as downstream enzymes phosphoglycerate mutase 2 (PGAM2) and enolase 3 (ENO3) catalyzing conversion to pyruvate all were significantly elevated in +IGF myotubes (Figure 8 B). We also looked at the involved voltage sensor calcium voltage-gated channel subunit alpha1 s (CACNA1S) that triggers Ca^2+^ release from the sarcoplasmic reticulum via the calcium channel ryanodine receptor 1 (RYR1). While elevated CACNA1S protein abundance from day 5 in +IGF myotubes did not reach significance, RYR1 protein levels were significantly upregulated starting from day 5 in myotubes differentiated in the presence of IGF1 (Figure 8 C). In summary, the abundance of proteins relevant for energy metabolism and sarcoplasmic Ca^2+^ release suggest that myotubes differentiated in the presence of IGF1 are better suited to cater to the demands of a physiological contraction.

**Figure 8.**
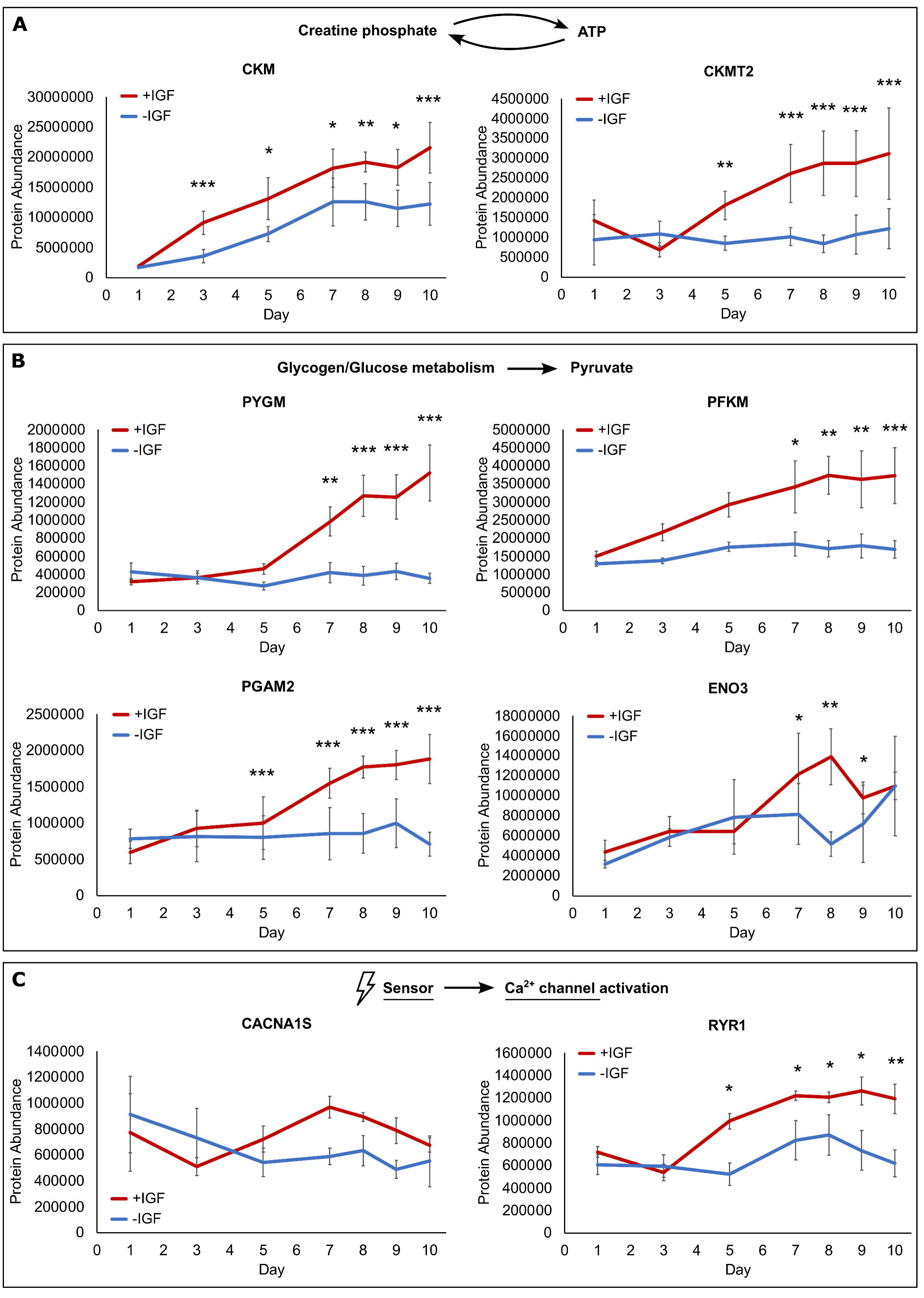
Energy metabolism and sarcoplasmic Ca^2+^ release proteins during human myotube differentiation. Primary human myotubes differentiated in the presence or absence of IGF1 were subjected to proteomic analysis over 10 days of differentiation. Regulation of proteins relevant for A) ATP supply (CKM, CKMT2), B) glycogenolysis and glycolysis (PYGM, PFKM, PGAM2, ENO3) and C) voltage dependent Ca^2+^ release (CACNA1S, RYR1) was analyzed on days 1,3, 5, 7-10 comparing protein levels between myotubes differentiated with or without IGF1. Curves represent mean ± SD, based on median values over all detected peptides. Significant differences were defined by a Benjamini-Hochberg corrected p-value below 0.05, *p < 0.05, **p < 0.01, ***p < 0.001, n = 4 individual donors.

### 4.8. Elevated Mitochondrial Protein Expression and Mitochondrial Respiration in +IGF Myotubes

Since the new differentiation protocol utilizing IGF1 supports differentiation of myotubes presenting more oxidative type I markers we investigated expression of all mitochondrial proteins comparing +IGF vs −IGF. Of the differentially expressed mitochondrial proteins between +IGF and −IGF at any stage of the differentiation (day 1-10), two thirds were upregulated in myotubes differentiated with IGF1 (heatmap left panel) and many of those were upregulated during differentiation compared to day 1 (heatmap right panels) (Figure S3 A). Among the upregulated mitochondrial proteins are subunits of the respiratory chain complexes (Figure S3 B).

Both, fiber type markers and mitochondrial proteome signature hint toward differentiation of more oxidative fiber type myotubes. Well in line, expression of PGC1α (*PPARGC1A*) RNA, a key regulator of mitochondrial biogenesis, maintenance and respiration, was strongly upregulated and elevated in myotubes differentiated with IGF1 starting from day 5 with a peak at day 8 (Figure 9 A). Thus, we next employed seahorse analysis to measure oxygen consumption and mitochondrial respiration (Figure 9 B-D). At all stages of the measurement, oxygen consumption rate (OCR) was elevated +IGF which was significant at baseline and with FCCP (Figure 9 B). The area under the curve over the complete measurement showed significantly elevated respiration in myotubes with IGF1 (Figure 9 C). Analyzing the respiratory states in detail, we found not only significantly elevated basal respiration but also more ATP production and uncoupled respiration as well as more spare capacity in myotubes differentiated in the presence of IGF1 (Figure 9 D). The energy map showed a shift from a quiescent to a metabolically active state in myotubes +IGF (Figure 9 E). The significantly elevated OCR/ECAR quotient indicates that this more driven by the increase in mitochondrial respiration than in glycolysis (Figure 9 E). In summary, in line with more oxidative fiber type markers and mitochondrial protein expression, myotubes differentiated in the presence of IGF1 do not only contract in a functional manner but show elevated mitochondrial respiration.

**Figure 9.**
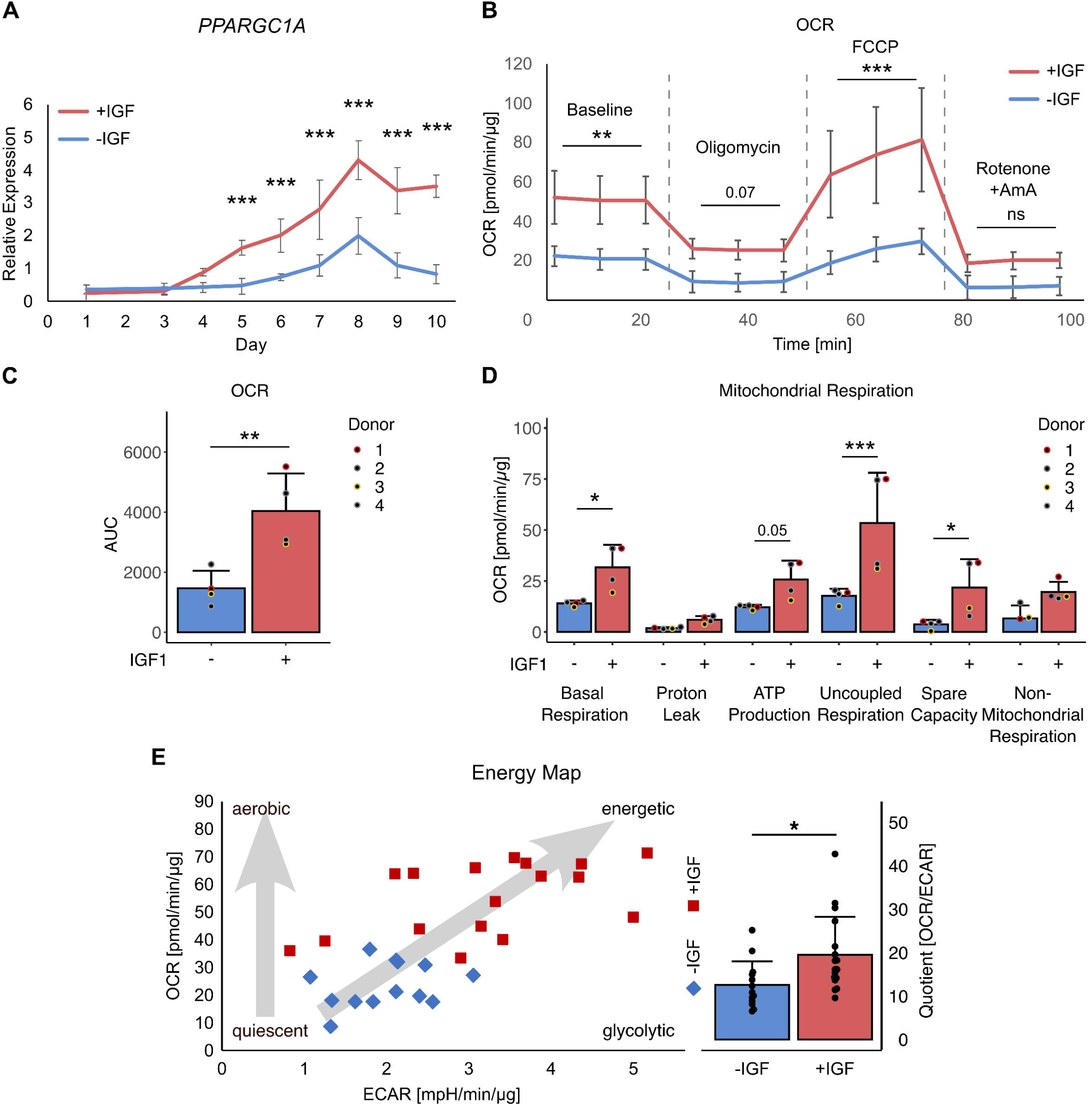
Mitochondrial respiration of human myotubes. Primary human myotubes were differentiated in the presence or absence of IGF1 over 10 days. A) RNA expression of *PPARGC1A* was analyzed on days 1-10 comparing expression between myotubes differentiated with or without IGF1. Curves represent mean ± SD. B) Respiration was measured in myotubes differentiated with and without IGF1 on day 8 of differentiation in response to indicated substrates using seahorse analysis. Lines represent mean ± SD of all analysis, labels describe OCR measurements after single injections indicated by the dashed lines. For statistical analysis all measurements between injections were compared. C) Area under the curve was calculated for the complete measurements and D) individual parameters of mitochondrial respiration calculated. E) Using baseline measurement at the third timepoint, the energy map was drawn relating OCR and ECAR. The quotient was calculated and compared +IGF vs −IGF. Significant differences were assessed using one-way ANOVA with Fisher’s LSD post hoc test, *p < 0.05, **p < 0.01, ***p < 0.001, n = 4 individual donors.

### 4.9. Insulin-Dependent Glucose Uptake in +IGF Myotubes

One important caveat of primary human myotubes *in vitro* as a model for diabetes research is the lack of proper expression of the insulin-dependent glucose transporter GLUT4. Therefore, we first looked at RNA expression of the relevant glucose transporters GLUT4 and GLUT1. Expression of *GLUT4* RNA was significantly elevated in myotubes with IGF1 starting at day 5 of differentiation (Figure 10 A). On protein level, the same trend was observed with significantly elevated GLUT4 protein on day 8 and 9 of differentiation in the presence of IGF1 (Figure 10 B). In contrast, GLUT4 RNA and protein expression stayed low in myotubes differentiated without IGF1 (Figure 10 B). Expression of GLUT1 RNA and protein was similar between myotubes +IGF and −IGF (Figure 10 C, Figure S4 A). Glucose uptake using a 2-deoxyglucose assay was measured after withdrawal of IGF for 48h, since the response to insulin shown as phosphorylation of AKT on serine-473 was impaired in myotubes that received IGF1 until they were stimulated with insulin (0h, Figure S4 B). Fasting IGF1 for 48h or 96h recovered the insulin sensitivity of +IGF myotubes (48h, 96h, Figure S4 B). As expected from the comparable GLUT1 abundance (Figure 10 A, Figure S4), both +IGF and −IGF myotubes showed similar basal glucose uptake in the 2-deoxyglucose assay (Figure 10 C). While myotubes differentiated without IGF1 were incapable of elevating glucose uptake after insulin stimulation, myotubes differentiated with the new protocol in the presence of IGF1 showed a 2-fold induction of glucose uptake after insulin stimulation. In summary, myotubes differentiated in the presence of IGF1 show functional contraction in line with elevated proper contractile apparatus protein assembly, differentiation of more oxidative fiber type myotubes in line with elevated mitochondrial respiration and elevated GLUT4 protein in line with elevated insulin-dependent glucose uptake.

**Figure 10.**
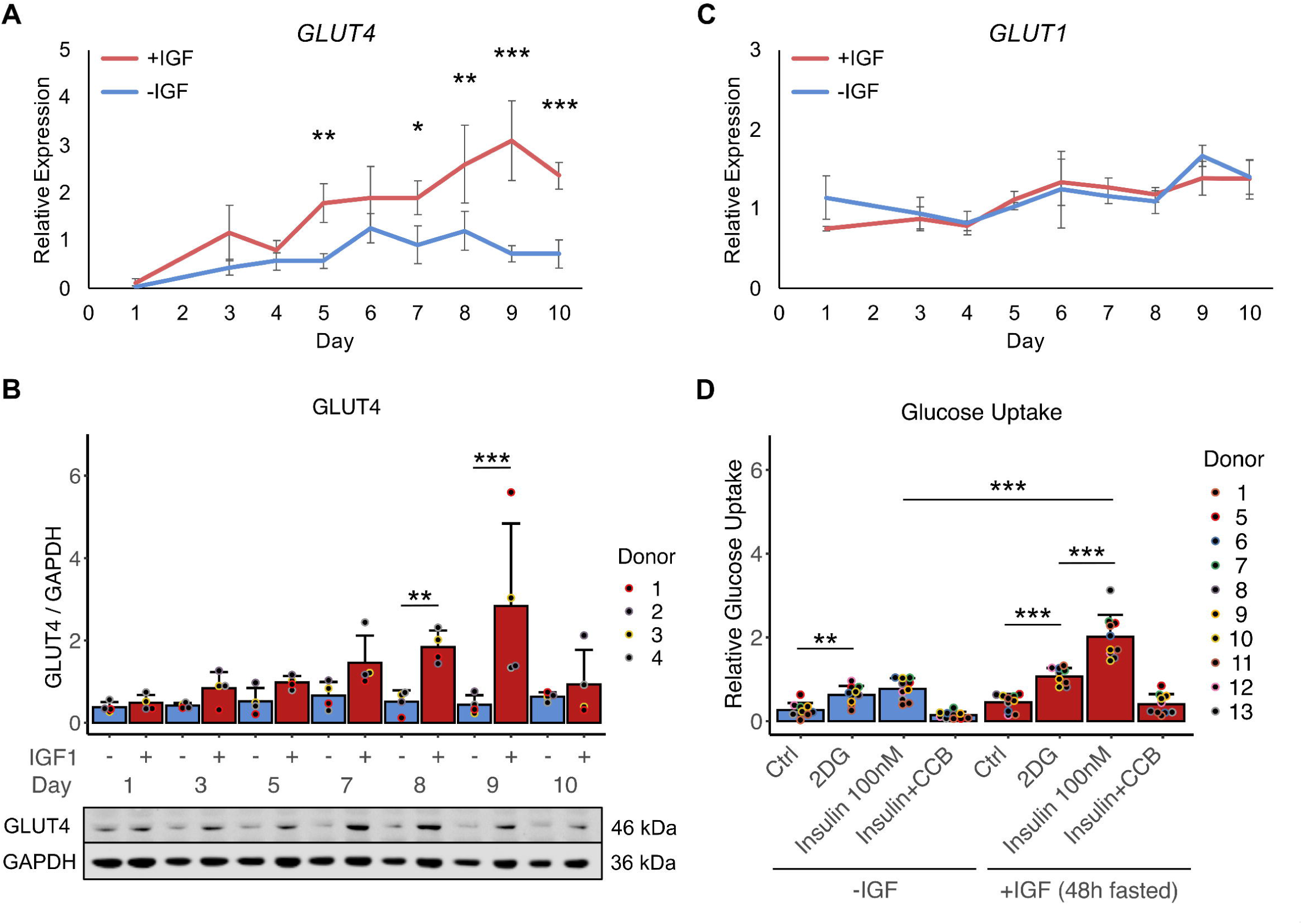
Glucose uptake of human myotubes. Primary human myotubes were differentiated in the presence or absence of IGF1 over 10 days. A) RNA expression of *GLUT4* and B) GLUT4 protein levels by western blotting were analyzed on days 1-10 comparing myotubes differentiated with or without IGF1. C) *GLUT1* RNA expression was analyzed +IGF vs −IGF over 10 days. Curves and bars represent mean ± SD, n = 4 individual donors. D) Glucose uptake, utilizing 2-deoxyglucose, was measured at baseline and in response to insulin stimulation in myotubes differentiated in the presence or absence of IGF1 on day 8. Bars represent mean ± SD, individual data points are depicted, n = 10 individual donors. Significant differences were assessed using one-way ANOVA with Fisher’s LSD post hoc test, *p < 0.05, **p < 0.01, ***p < 0.001.

## 5. Discussion

In this study, we developed and characterized a culture system to achieve functionally active, contractile human myotubes suitable to study metabolic behavior and parameters relevant for exercise and diabetes research. Based on a serum-free system, the sole addition of IGF1 drives the cells toward more mature differentiated myotubes capable of contraction, insulin-dependent glucose uptake and oxidative phosphorylation. We not only functionally validated the phenotype but demonstrated a pronounced shift in the proteome compared to myotubes differentiated without IGF1, which is responsible for the improved functionality.

Many growth factors are important for muscular development and myotube differentiation. Myotube growth, differentiation and hypertrophy are primarily regulated by a signaling pathway initiated by insulin-like growth factor 1 (IGF1) [45]. *In vivo*, IGF1 can be secreted by myofibers, macrophages and endothelial cells in response to injury and exercise, or reach muscles through the circulatory system inducing an intracellular cascade after binding its receptor [46; 47]. IGF1 induces Akt signaling which via mTOR promotes protein synthesis and hypertrophy while inhibiting protein degradation and muscle atrophy by impeding GSK3 and the FOXO family of transcriptions factors [48; 49]. The mTOR signaling pathway also plays a key role in skeletal muscle differentiation by upregulating expression of Pax7, Myf5, MyoD, and myogenin [50]. Besides IGF1, HGF, FGF, and PDGF have been associated with muscle repair and hypertrophy by activation of satellite cells [51]. However, all three, HGF, FGF, and PDGF had been shown to inhibit myogenic differentiation maintaining a proliferative phenotype of satellite cells and myoblasts [47; 52]. Thus, for a growth factor to improve myotube differentiation we focused on IGF1 in this study.

IGF1 was previously investigated in human myotubes with regards to delaying or overcoming myoblast senescence and to induce hypertrophy with singular IGF1 doses or 1h treatments [33; 53; 54]. In these studies, IGF1 treatment was already described to increase myotube diameters and helped to overcome the detrimental effects of cotreatment with TNF-α on myotube differentiation [54]. Some studies focused on a specific isoform of IGF1 termed mechano-growth-factor or MGF (IGF1c in human, IGF1b in rodents) that is only detectable in tissue after mechanical strain and/or damage [55; 56]. MGF was suggested to be very effective in promoting myotube differentiation in one study [57] only to be found less effective than recombinant IGF1 by another study reporting stronger improvement in myotube area and diameters with IGF1 than MGF [58]. In this study, we further improved primary human myotube differentiation by supplementing human recombinant IGF1 in a serum- and insulin-free approach and extensively characterized the functional and metabolic phenotype of differentiated myotubes beyond the previously described benefits on diameter and hypertrophy.

Mimicking exercise in *in vitro* cultures utilizing EPS with primary human myotubes yields molecular responses that are relatable to *in vivo* responses of skeletal muscle to exercise [59; 60]. Often these studies fail to report a visually contracting phenotype or if provided, show diminutive degrees of movement likely due to the limitations of previous *in vitro* differentiation protocols and the use of serum [27; 28]. In a previous study with the initial focus on co-culture of human myotubes with motoneurons Guo et al. used a labor-intensive protocol with several distinct growth factor cocktails in subsequent use during primary human myotube culture [25]. They focused on electrophysiological properties of the differentiated myotubes and reported spontaneous contractions in culture as well as EPS-induced contraction comparable to our myotubes. Being a vastly more complex approach, their protocol contained two similarities to our approach. Apart from containing a mix of neurotrophins (BDNF, GDNF, NT3/4, CNTF) and other additives their differentiation media 2 and 3 contained no serum and a small amount of IGF1 (10 ng/ml). We also previously described the beneficial effects of using completely serum-free myotube differentiation with respect to the myosin heavy chain expression profile, PGC1α (*PPARGC1A*) expression levels and number of non-fused cells already without IGF1 supplementation [24]. By adding only IGF1 to our previous “easy to use” serum-free differentiation protocol we achieved a comparable degree of functional myotube differentiation already on day 8 as previously reported for day 10 [25]. Furthermore, our proteomic analysis provides a comprehensive characterization and molecular basis for the capability to contract by revealing elevated levels of contractile apparatus proteins as well as proteins involved in energy metabolism such as ATP-supply, glycogenolysis, glycolysis and voltage dependent Ca^2+^ release from the sarcoplasm, essential for muscular contraction. *In vitro*, primary human myotubes are generally considered to have a lower mitochondrial oxidative capacity preferring glycolysis over lipid oxidation [26]. Accordingly, human myotubes show elevated MYH1 and MYH2 expression over MYH7 which is characteristic for the glycolytic type II fiber type [26]. We here found that IGF1-guided myotube differentiation led to upregulation of slow type I MYH7, MYH6 and MYL3 proteins suggesting that IGF1 drives myotube differentiation toward a more oxidative phenotype. Animal studies demonstrated the importance of PGC1α for fiber-type switching from glycolytic fast type to oxidative slow type muscle fibers [61; 62]. Thus, the robust upregulation of PGC1α provides one potential explanation for the increased oxidative capacity of IGF1-treated myotubes, which was finally validated by analyzing mitochondrial respiration. Altogether this data convincingly shows that IGF1-guided differentiation of human myotubes overcomes the glycolytic fiber type bias by shifting differentiation toward more oxidative slow type I fiber type myotubes.

Another limitation of human myotubes *in vitro* was that, compared to *in vivo*, the insulin-stimulated glucose uptake appears impaired, accompanied by reduced GLUT4 expression at least in relation to GLUT1 [26]. *In vivo*, GLUT4 is more highly expressed in type I oxidative muscle fibers that are generally regarded as having a greater insulin-stimulated glucose uptake [63; 64]. Our IGF1-guided differentiation promoted development of more oxidative type I myotubes and in line, we found elevated GLUT4 mRNA and protein expression at similar GLUT1 levels. Because of the high similarity of insulin- and IGF1-receptors, the existence of hybrid receptors and the potential of IGF1 to bind to the insulin receptor although with lower affinity, combined with a substantial overlap of downstream signaling via IRS and AKT, it is possible that cells treated with IGF1 can become insulin resistant [65; 66]. In line we saw reduced responsiveness to 10 nM insulin stimulation for 10 min in AKT phosphorylation in myotubes cultured with continuous IGF1 supplementation compared to 48h or 96h of IGF1 fasting. After 48h of IGF1 fasting, myotubes differentiated with IGF1 where not only insulin responsive but also displayed a 2-fold elevated glucose uptake in response to insulin stimulation that myotubes differentiated without IGF1 were incapable of. Therefore, we were able to show that IGF1-guided myotube differentiation is able to improve the shortcomings of previous differentiation protocols regarding GLUT4 expression and glucose uptake.

We assume that the pronounced effect on functional myotube differentiation with our protocol can be attributed to the combination of a serum-free basal medium with IGF1 supplementation. The skeletal muscle- and contractile apparatus-specific protein expression is in line with the role of IGF1 in skeletal muscle development. The overall improvements compared to previous studies, sometimes also utilizing IGF1, are however likely observed due to a more unobstructed IGF1 activity in serum-free differentiation medium. Serum can contain other growth factors such as WNT, TGF-β or BMP agonists, that can induce antagonizing signaling pathways inhibiting the beneficial effects of IGF1 signaling, or IGF-binding proteins that directly bind and inhibit IGF1 [67; 68]. Thus, our data show and give suggestions why IGF1-guided human myotube differentiation overcomes the previously inadequate capability of established human myotube models with our newly described “easy to use” differentiation protocol.

We are aware that high levels of functionality can also be achieved and observed with 3D culture tissue engineering approaches also giving way to even more elaborate readouts [69–71]. However, it was our goal to develop an “easy to use” 2D protocol which can be implemented in many laboratories, driven by metabolic or diabetes related research questions, where monolayer myotube culture is readily available while establishing complicated tissue engineered 3D solutions is no valid option. For these use cases our protocol allows studies with a more functional human myotube with the same culture methods most researchers are familiar with.

As we already observed that IGF1 does not need to be present for the last 48h of differentiation to still yield benefits of IGF1-guided human myotube differentiation regarding insulin-dependent glucose uptake it is possible that this protocol can be further optimized by reducing the time period that IGF1 is given to stimulate proper myotube differentiation as effects on myotube diameter have been reported for singular treatments in serum containing media before [54; 58]. Also, it is possible that beneficial effects can be achieved with lower concentrations as previously also lower concentrations were used to report some effects on hypertrophy [33; 58]. Thus, lower concentrations or shorter periods of IGF1 treatment might be sufficient for the reported beneficial effects on differentiation. In future projects, our new differentiation protocol can be employed to generate a microfluidic, inter-chip compatible human exercise-on-chip model. This is in particular of relevance as for many exercise-related research questions, the release of myokines and interorgan crosstalk, novel culture techniques like organ-on-chip are favorable.

One limitation of this extensive characterization of an IGF1-guided human myotube differentiation is that the time course data due to handling limitations could only be performed in parallel in 4 donors that were all overweight. However, age and weight of the donors projected a substantial range and 2 males and 2 females were chosen. As qRT-PCR and proteomics data presented with robust results and low variation, and most functional analysis were repeated in additional donors we consider the dataset robust and appropriate to underline our conclusions.

## 6. Conclusion

Utilizing IGF1 in a serum-free differentiation medium, we here describe a novel “easy to use” differentiation protocol for functional primary human myotubes from isolated satellite cells. Compared to previous published protocols this approach can be easily adopted by anyone working with human myoblasts. The obtained myotubes within a week of differentiation recapitulate the physiological traits of skeletal muscle *in vivo* vastly superior to established 2D *in vitro* protocols. We report improvements in contractile movement, capability for excitation, glycogen and glucose metabolism to cater to the energy demand of a physiological movement of myotubes, mitochondrial respiration and insulin-dependent glucose uptake. Especially as a model for diabetes and exercise research, the capability to contract together with a more oxidative phenotype open up novel possibilities to study the molecular processes underlying the beneficial effects of exercise to prevent the onset of type 2 diabetes. Our results are also valuable for tissue engineering of linearly aligned myobundles in perfusable bioreactors and transfer to microfluidic systems for an exercise-on-chip model for drug screening and development of an interorgan crosstalk model.

## 8. Acknowledgements

The authors thank all biopsy donors. The authors are grateful for the excellent technical support provided by Nadine Vilas from the University Hospital Tübingen, Tübingen, Germany.

## 11. Author contributions

S.I.D., C.W., P.G. conceived and designed research; P.G., C.v.T., A.M., J.M. performed experiments; S.I.D, C.v.T., S.H., T.G. analyzed data; S.I.D, C.v.T., C.W. interpreted results of experiments; S.I.D prepared figures; S.I.D., C.W. drafted manuscript; S.I.D, C.W. edited and revised manuscript; S.I.D., P.G., C.v.T., A.M., J.M., T.G., A.L.B., A.P., P.L., S.H., C.W. contributed to discussion and approved final version of the manuscript.

## 12. Data Availability

The proteomic data used in this study will be available via PRIDE - Proteomics Identification Database Project accession: PXD043200. Further data that support the findings of this study are available from the corresponding author upon request.

---

Video 1

Human myotubes differentiated with IGF1 representative for best contraction at 10x magnification

Video 2

Human myotubes differentiated with IGF1 representative for best contraction at 10x magnification

Video 3

Human myotubes differentiated with IGF1 representative for average contraction at 10x magnification

Video 4

Human myotubes differentiated with IGF1 representative for average contraction at 10x magnification

Video 5

Human myotubes differentiated with IGF1 representative for worst contraction at 10x magnification

Video 6

Human myotubes differentiated with IGF1 representative for worst contraction at 10x magnification

Video 7

Human myotubes differentiated with IGF1 and visible striation at 30x magnification

Video 8

Human myotubes differentiated without IGF1 representative for average at 10x magnification

Video 9

Human myotubes differentiated without IGF1 representative for average at 10x magnification

Video 10

Human myotubes differentiated without IGF1 representative for best at 10x magnification

Video 11

Human myotubes differentiated with IGF1 showing spontaneous contraction at 10x magnification

Video 12

Human myotubes differentiated with IGF1 showing spontaneous contraction at 10x magnification

Video 13

Human myotubes differentiated with IGF1 spontaneously contracting followed by EPS at 10x magnification

Video 14

Human myotubes differentiated with IGF1 showing tetanic contraction at 30x magnification

