## Supplementary Figures for "Engineering of human myotubes toward a mature metabolic and contractile phenotype"

### 1. Supplementary Figures

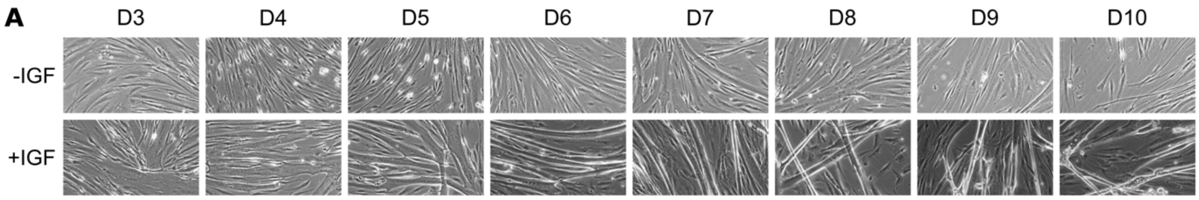


Figure S 1

Microscopic evaluation of human myotube differentiation

Primary human myotubes were differentiated in the presence or absence of IGF1 over 10 days. A) Representative microscopic images of differentiated myotubes from day 3 to day 10. All images were taken at 20x magnification and were not cropped.

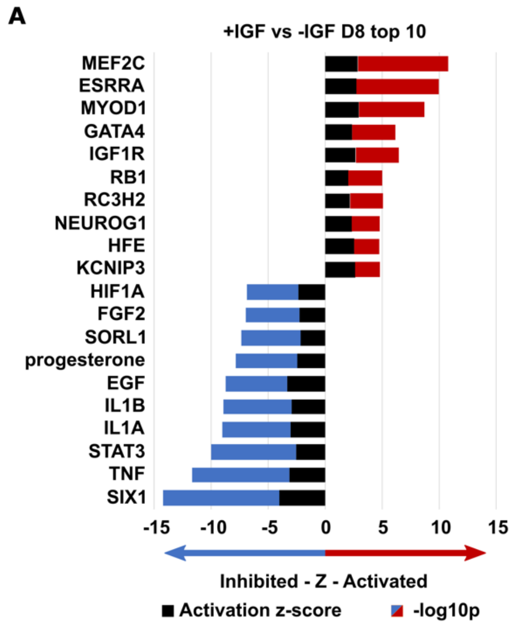


Figure S 2

Ingenuity Upstream analysis +IGF vs -IGF

Primary human myotubes were differentiated in the presence or absence of IGF1 for 8 days. A) Ingenuity upstream analysis was performed based on significantly differentially regulated proteins comparing +IGF vs -IGF n=4 individual donors. Stacked bars represent z-score and -log10 p value for activated (red) and inhibited (blue) upstream regulator pathways.

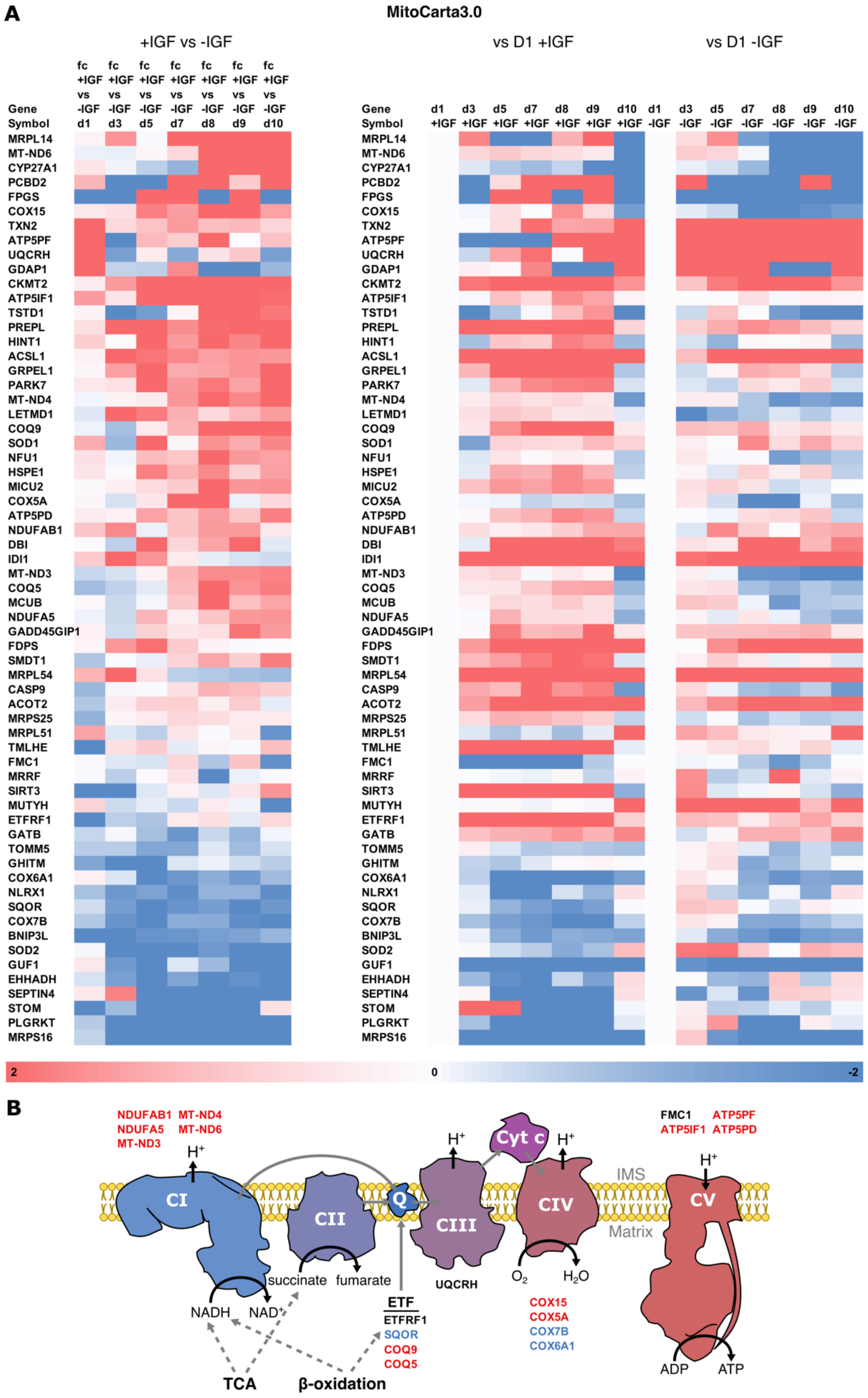


Figure S 3

Mitochondrial proteins during human myotube differentiation

Primary human myotubes were differentiated in the presence or absence of IGF1 over 10 days. A) Heat map of all significantly regulated proteins associated to mitochondria based on MitoCarta3.0. The left panel shows differential regulation comparing +IGF vs -IGF. Right panels show regulation compared to day 1 in +IGF (left) and -IGF (right). Colors represent fold changes, significant differences were defined by a Benjamini-Hochberg corrected p-value below 0.05, *p < 0.05, **p < 0.01, ***p < 0.001, n = 4 individual donors. B) Schematic representation of regulated proteins of the respiratory chain generated with InkScape (v1.0), colors indicate direction of regulation in +IGF vs -IGF.

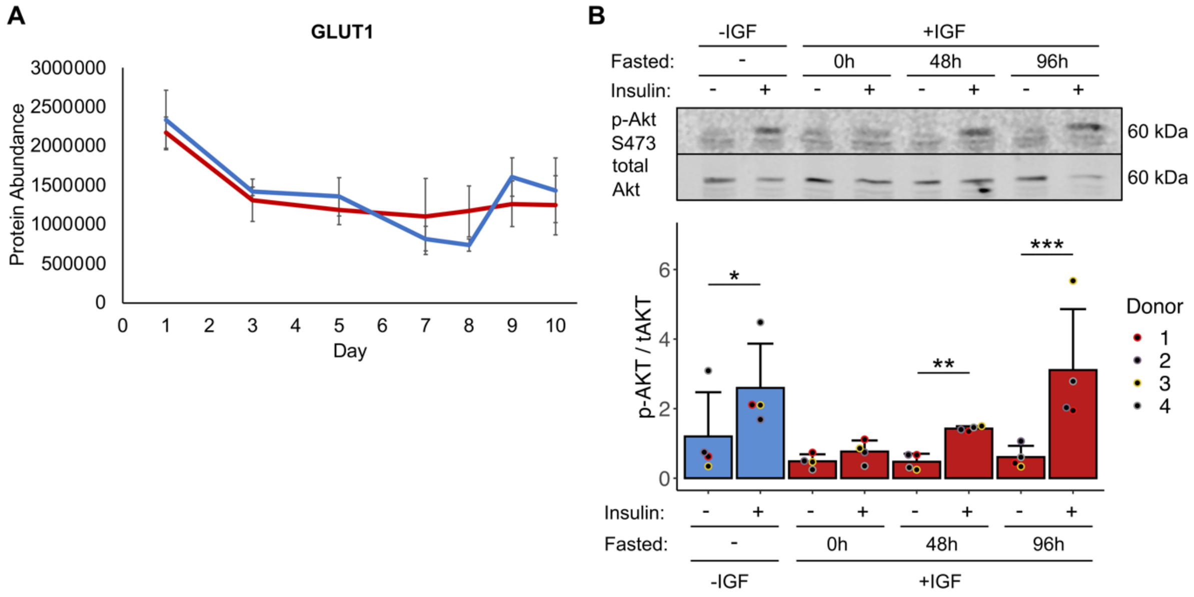


Figure S 4

GLUT1 protein during human myotube differentiation and insulin responsiveness

Primary human myotubes differentiated in the presence or absence of IGF1 were subjected to proteomic analysis over 10 days of differentiation. A) Regulation of GLUT1 protein abundance was analyzed on days 1,3, 5, 7-10 comparing protein levels between myotubes differentiated with or without IGF1. Curves represent mean ± SD, based on median values over all detected peptides. n = 4 individual donors. B) Phosphorylation of AKT (S473) was analyzed in response to 10nM insulin stimulation for 10 min in myotubes differentiated without IGF1 and myotubes differentiated with IGF1 either without, with 48h or 96h of fasting IGF1 before harvest on day 8 of differentiation. Bars represent mean ± SD, individual data points are depicted, n = 4 individual donors. Significant differences were assessed using one-way ANOVA with Fisher's LSD post hoc test, *p < 0.05, **p < 0.01, ***p < 0.001.
